# Plant uses in a traditional fisherman community in northeastern Brazil

**DOI:** 10.1101/620542

**Authors:** DYP Tng, DMG Apgaua, MDS Lisboa, CN El-Hani

**Affiliations:** Instituto de Biologia, Universidade Federal da Bahia, National Institute of Science & Technology in Interdisciplinary and Transdisciplinary Studies in Ecology and Evolution (INCT IN-TREE), Brazil. R. Barão Jeremoabo, Ondina, 40170-115 Salvador, Bahia, Brazil; Centre for Tropical, Environmental and Sustainability Sciences, College of Science and Engineering, James Cook University, 14-88 McGregor Rd, Smithfield, Queensland 4878, Australia

**Keywords:** artisanal fishermen, Bahia, northeast Brazil, traditional plant knowledge

## Abstract

**Background:** Knowledge of traditional plant use in rural communities is under threat from urbanization and also land use change. Hence, elucidating region-specific patterns traditional knowledge of habitat resource use is crucial for assisting policy making with regard to nature conservation, human nutrition, and human health. Using original data from northeast Bahia, Brazil, we aim to document the collective knowledge of plant use possessed by artisanal fishermen and women in fishing communities, related to medicinal, construction, technology and ritualistic/religious uses.

**Methods:** Data were collected through interviews with and opportunistic guided tours by local specialists to record the local knowledge of plant use and methods of use.

**Results:** Through interviews, 116 species of plants from 104 genera and 47 botanical families were identified along with their local names, plant parts utilized, habitats, and preparation methods, and an additional 26 spp. for which only local names were available. Of these, 72 spp. are used as medicine, 48 spp. as non-conventional edibles, 31 spp. for construction and 11 for religious/mystical purposes. Female informants generally cited more food and medicinal plants than male informants. All interviewees unanimously expressed that the use of plants for medicines and fishing technology has significantly reduced or been superseded by modern materials.

**Conclusion:** The present study demonstrates that the rich collective plant use knowledge of an artisanal fishing community in northeast Brazil. The results of this study serves as a framework for to extend ethnobotanical investigations to other traditional fishing communities in the vicinity, and also to examine more social and demographic factors influencing traditional knowledge related to uses of plants. Finally, the rich tradition of plant use in the region underpins the need to encourage measures to preserve this cultural knowledge and conserve the natural environments that are a source of these plants.

## INTRODUCTION

Throughout history, wherever humans have settled into communities they have been faced with the necessity of developing strategies for managing and using natural resources to meet their daily needs. The unique cultural heritage within any given traditional community is thus comprised in part by the culmination of knowledge of the local environmental and the practices associated with uses of natural plant, animal and geological resources. However, the erosion of culture in traditional communities worldwide is occurring at an alarming and unprecedented rate due to modernization (Benz et al. 2000; Tang & Gavin 2016) and also by the development of tourism (Briassoulis & van der Straaten 2013; Canavan 2016; Xue et al. 2017).

In terms of biodiversity conservation, several authors have expounded on the importance of conserving ethnobotanical knowledge as part of living cultural knowledge and practice between communities and the environment for biodiversity conservation (Maroyi 2011). Research on local knowledge also confirm the value of these cultures and self-sufficiency of local communities, as well as the importance of biodiversity to the sustainability of these populations (di Stasi et al. 2002).

Communities that occupy areas close to natural vegetation such as tropical forests coexist with great biological diversity, and possess a cultural repertoire of practices related to the uses of plants in everyday life (de Santana et al. 2016; Sujarwo et al. 2016). In Brazil, these communities include indigenous ethnic people, Quilombolas (descendants of Afro-Brazilian runaway slaves) and various traditional communities of fishermen, river-dwellers, rubber sap gatherers, etc, who often live in well-delineated communities in rural areas away from major cities and surrounded by natural landscapes.

Medicinal plants play a hugely important role in the well-being of these rural communities, and to improve the use of the Brazilian biodiversity and public access to herbal medicines the Brazilian government has encouraged the use of medicinal plants and herbal medicines (Brasil 2006). Indeed, among the various categories of traditional plant uses, the use of medicinal plants is by far the most well studied. In Northeastern Brazil for instance, there have been various ethnobotanical studies focused on medicinal plant use in various traditional communities (Alberquerque et al. 2008; Borges & Bautista 2010; Cartaxo et al. 2010; Cunha et al. 2012; da Silva et al. 2012; Gomes & Bandeira 2012; de Araújo et al. 2018). In spite of these surveys, the geographical coverage of studies on plant use in traditional communities is still far from complete. For instance, there are still very little studies on communities in the littoral northeast of Bahia, where traditional communities that practice artisanal fishing still have a rather well preserved culture.

As a corollary to medicinal plant use, ethnic indigenous communities often have systems of religious or ritual plant use for healing or for restoring well-being (Bussman & Sharon 2006) the use of which is poorly documented in the region (Almeida et al. 2014). In these instances, there are members of the community who serve in the capacity of shamans or spiritual healers who help cure ailments through ritual use of plants (). By comparison, such practices are less well known in traditional communities (Rodrigues & Carlini 2006), but at the same little has been documented about any ritual use of plants in such communities.

Additionally, other aspects of plant use deserving attention include the use of non-conventional plant foods, here broadly defined as wild, wayside, or naturalized plants that can supplement the nutrition of people in rural communities (Barminas et al. 1998; Kebu & Fassil 2006). Interest in non-conventional plant foods in Brazil have increased in recent years (Kinupp & Lorenzi 2014; do Nascimento et al. 2015) but this aspect of traditional plant use remains poorly studied in most regions in Brazil.

Perhaps one of the most poorly documented aspect of traditional plant uses are those used for construction, tool making and technology. This may be particularly relevant for traditional communities who rely on fishing as the main source of sustenance, as fishermen would have utilized plant resources to construct boats, and other structures and tools involved in their fishing technology. Focusing on a traditional fishing community in the northeast of Brazil, we take a holistic approach in our objective to document traditional knowledge of plants for medicine, food, construction and religious use. As gender has been widely studied as a potential driver of plant knowledge (Torres-Avilez et al. 2016), we also compare the patterns of plant knowledge held by male and females in the community, with the hypothesis that there will be a divergence in male and female knowledge about plant use.

## MATERIAL AND METHOD

### Study area and historical context

The study was conducted in Siribinha (11°48′49′S, 37°36′38′W), an artisanal fishing community with a population of ∼400 located in proximity to the mouth of the Itapicuru river on the north coast of the State of Bahia, Brazil. This community is within the Municipality of Conde, and has an economy driven primarily by tourism, coconut plantations, fishing, and the cattle farming.

The village of Conde was part of the Portuguese occupation north of Salvador, which began in 1549 with the first cattle corrals around the present city. Between 1558 and 1572, Jesuit missionaries arrived to the area where Conde is today, interacting with the indigenous Tupinambás indigenous people who occupied the region. In 1621, Portuguese settlers started to raise livestock and sugarcane and tobacco crops with slave labor, and in 1806, Conde became an area of litigation between the states of Bahia and Sergipe. By 1935, Conde was formally recognized as a municipality within the state of Bahia, with headquarters in the town of Ribeira do Conde.

The natural vegetation of the area consists of freshwater alluvial wetlands, mangroves, beach vegetation, shrubby thicket-like forests (locally-known as *restingas*) growing on sand dunes, and remnant patches of closed forest (Atlantic forest). Coconut plantations and cattle ranches also make up part of the land use tenure of the municipality.

Part of the territory of Conde is located within the Environmental Protection Area of the North Coast (http://www.oads.org.br/leis/2724.pdf), but this has not prevented socio-environmental problems such as deforestation and disorderly appropriation of territory which threatens the cultural heritage of the municipality and ecological integrity of surrounding natural habitats (e.g., Limonad 2007). In particular, residents in several coastal settlements in the Municipality of Conde face socio-economical problems of limited employment and income opportunities, leading to an exodus of the younger generation to larger cities, and thereby contributing to the erosion of traditional culture.

### Interviews and random walks with traditional experts

The study was conducted between August 2018 to February 2019. During this period, we made door-to door visits in order to identify local people with a specialized knowledge of plant use (Davis and Wagner 2003). Therefore, our sampling approach was intentionally non-random (Albuquerque et al. 2014a), under the assumption that local specialists would provide more specific and higher quality information concerning medicinal plants. During our first contacts with the communities the inhabitants themselves identified specialists, who we define as “individuals recognized by the community as having deep knowledge about the traditional uses of native and/or introduced plants”. Using the snowball method (Bailey 1994), names of other specialists were then obtained. Eventually, we interviewed a total of 21 local specialists (11 females and 10 males in Siribinha) were eventually interviewed, with ages ranging from 43-82. All the local specialists identified had lived in the community for at least 30 years, and were frequently sought out by other community members for advice on plant use. The men are currently active or retired fishermen, and the women consisted of home-makers who sometimes fished or specialized in collecting crabs (*marisqueras*). One of the female local specialist interviewed was also regarded in the community as a “*rezeira*”, or one who prays or bestows blessings.

We used semi-structured interviews (Albuquerque et al. 2014b), which best suited the preferences of the community inhabitants. All interviews were conducted in Brazilian Portuguese, and specialists were interviewed individually as recommended by Phillips and Gentry (1993), to ensure that responses were independent. As was appropriate for the moment, during the course of interviews some traditional experts brought us on guided tours (Albuquerque et al. 2014b) through their gardens or through the natural vegetation in the vicinity of their houses to describe the plants they use. Our questionnaires gathered information about the medicinal, food, construction and ritual uses of plants, plant names, plant parts used, and preparation methods where relevant.

As our aim was to evaluate how community members use plants into 12 categories: medicine, food, for implement-making for fishing practices or domestic use, construction, and ritual use, following modified categories of Prance et al. (1987). For the category of plants used as non-conventional food, we consider only species collected from the surrounding environments or planted in gardens to supplement conventional food. This includes non-conventional plant food species following the definition of Kinupp & Lorenzi (2014), and native or naturalized fruits harvested from surrounding habitats (Lorenzi et al. 2015). We also distinguished the lifeforms of the plants cited (tree, shrub, herb and climber).

Plants were identified in the field or where identification was not possible, a sample was brought back to the lab and identified at the herbarium at the Federal University of Bahia, using the literature (Lorenzi & Matos 2002) or through consultations with experts. Voucher specimens of all collected samples were lodged at the Laboratory of Teaching, Philosophy and History of Biology at the Federal University of Bahia. Plants were classified in terms of their origin as native (native to Brazil), or non-native and naturalized based on the online flora of Brasil (floradobrasil.jbrj.gov.br/2012/), and the cultivated status of each species was noted.

### Data analysis

The first step in our data analysis was to calculate two indices related to plant use, the use value (UV) and the informant concensus factor (ICF). The UV demonstrates the relative importance of each species known locally, which we calculated following the formula: UV = U/N, where U refers to the number of citations per species, and N refers to the number of traditional experts interviewed (Gürdal and Kültür 2013). The ICF denotes the degree to which traditional experts exchange information about plant use, which we calculated using the following formula: FIC = N_ur_ - N_t_/N_ur_ - 1, where Nur refers to the number of use citations in each category and Nt is the number of species indicated in each category (Cartaxo et al. 2010). Therefore, ICF values will be low (approaching 0) if plants are chosen randomly, or if traditional experts do not exchange information about their use, but high (approaching 1) if information is exchanged between them. For the ICF index, all citations were placed into 12 plant use, distribution type or lifeform categories: medicine, food in general, non-conventional food plants, construction and technology in general, fishing-related technology only, ritual and mystical plant use, native, non-native, tree, shrub, herb, and climber.

With the 21 traditional experts, we tested the difference in the number of plants cited by male and female traditional experts using a student’s t-test (α = 0.05). We also identified major gradients in plant use knowledge using non-metric multidimensional scaling (NMDS) ordination, using a Jaccard distance matrix. For this analysis we included all 142 cited plants, and performed Spearman-rank correlations for each species with each NMDS axis to examine the main species driving patterns in the ordination.

## RESULTS

We recorded 116 species belonging to 104 genera and 47 families (Table 1). The identities of an additional 26 cited plant records could not be ascertained and only local names are presented (Table 1). The top five botanical families with the most number of species with cited uses were the Fabaceae (14 spp.), Arecaceae, Lamiaceae and Myrtaceae (7 spp. each) and the Anacardiaceae (6 spp.) (Table 1). Plants with the highest Use Values (UV) were *Rhizophora mangle* and *Anacardium officinale* (both UV = 0.52), *Cocos nucifera* and *Myrciaria floribunda* (both UV = 0.48), and *Manilkara salzmanii* (both UV = 0.43).

**Table 1.**
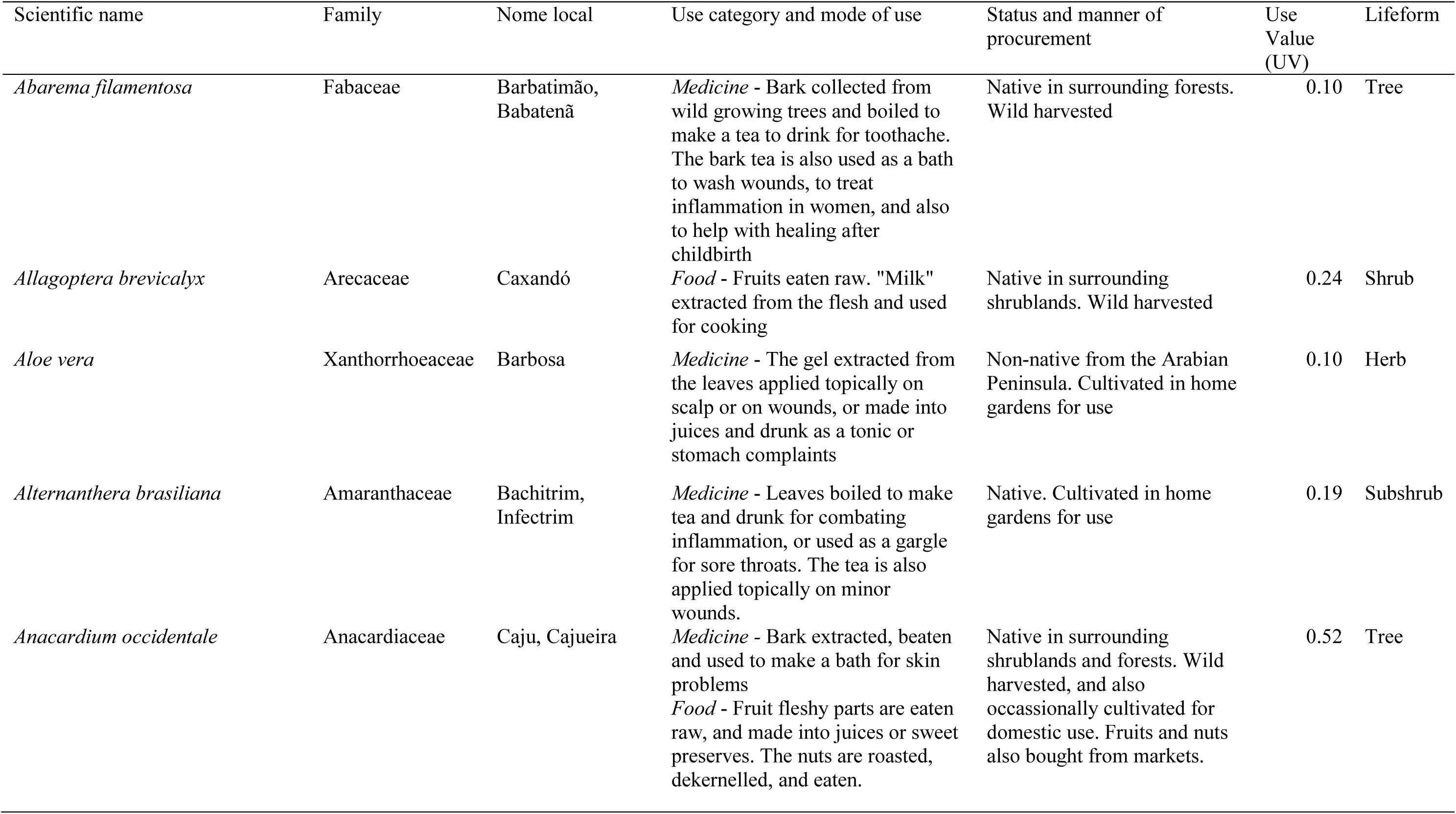
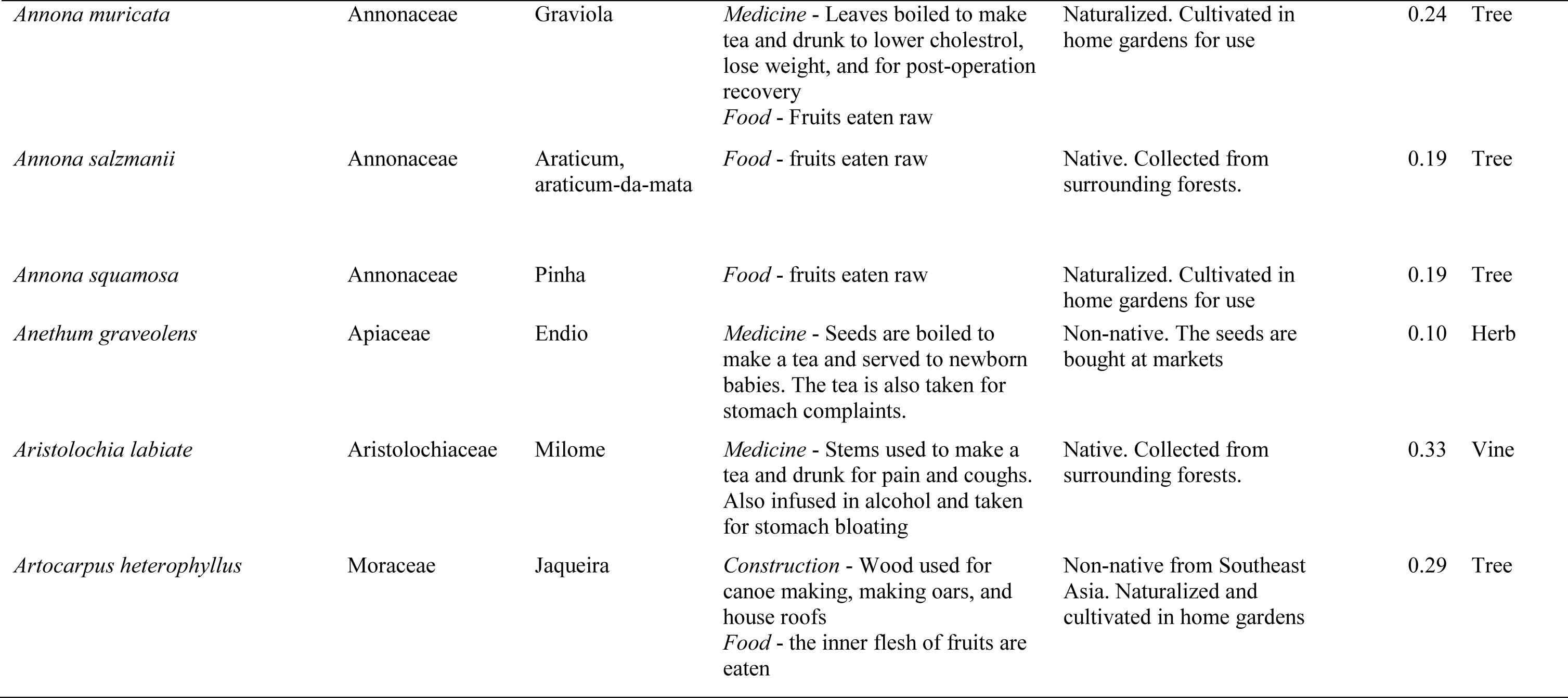
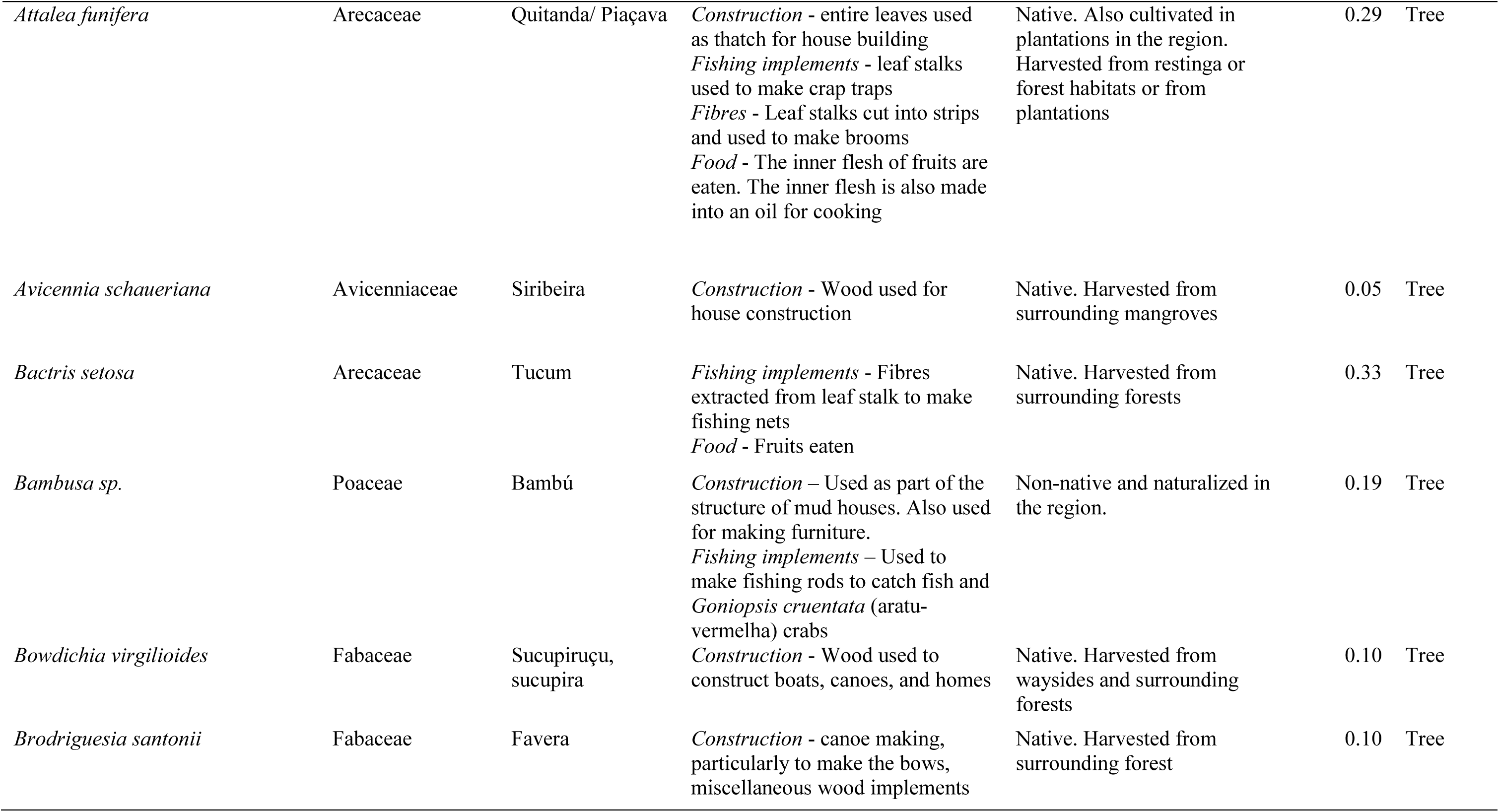
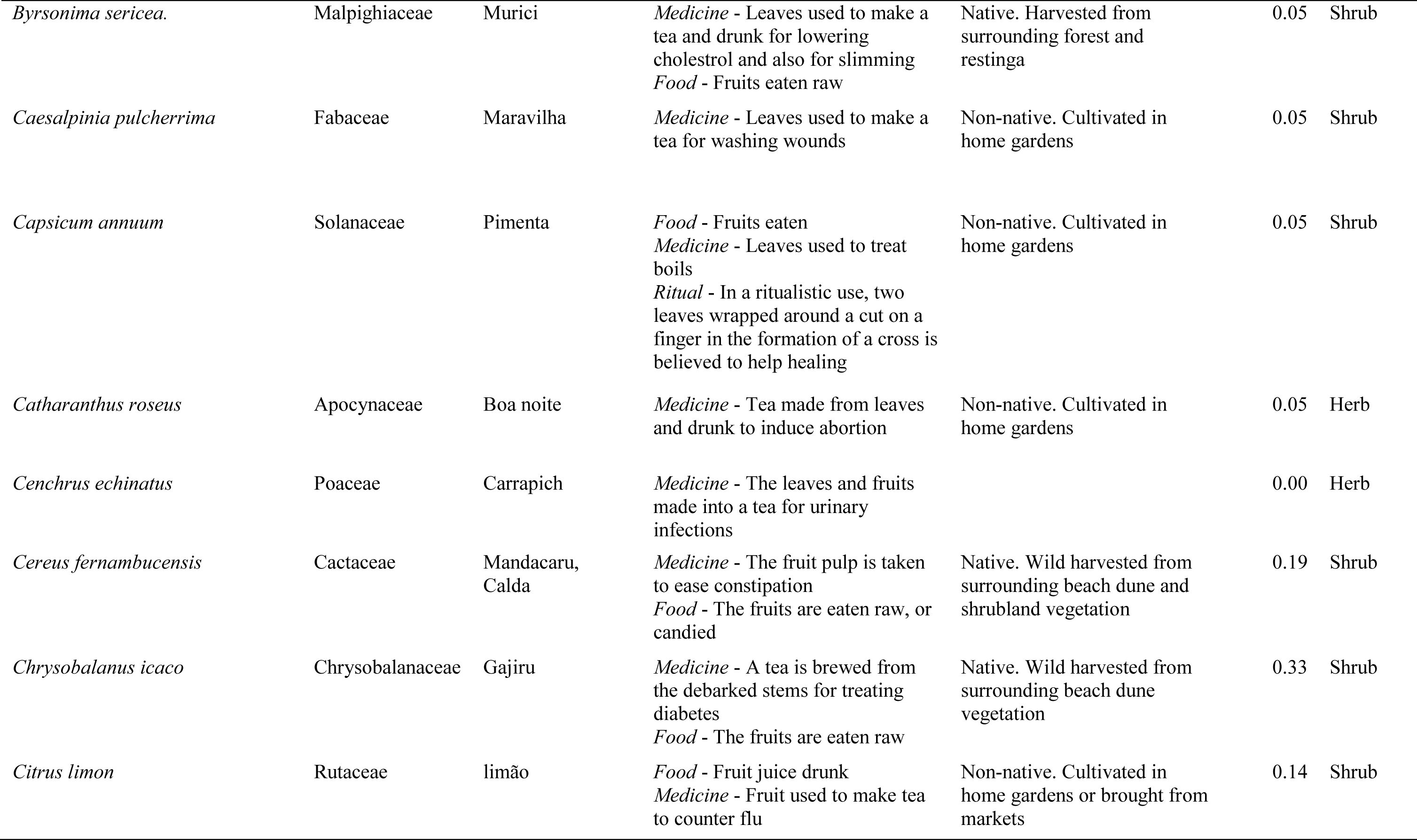
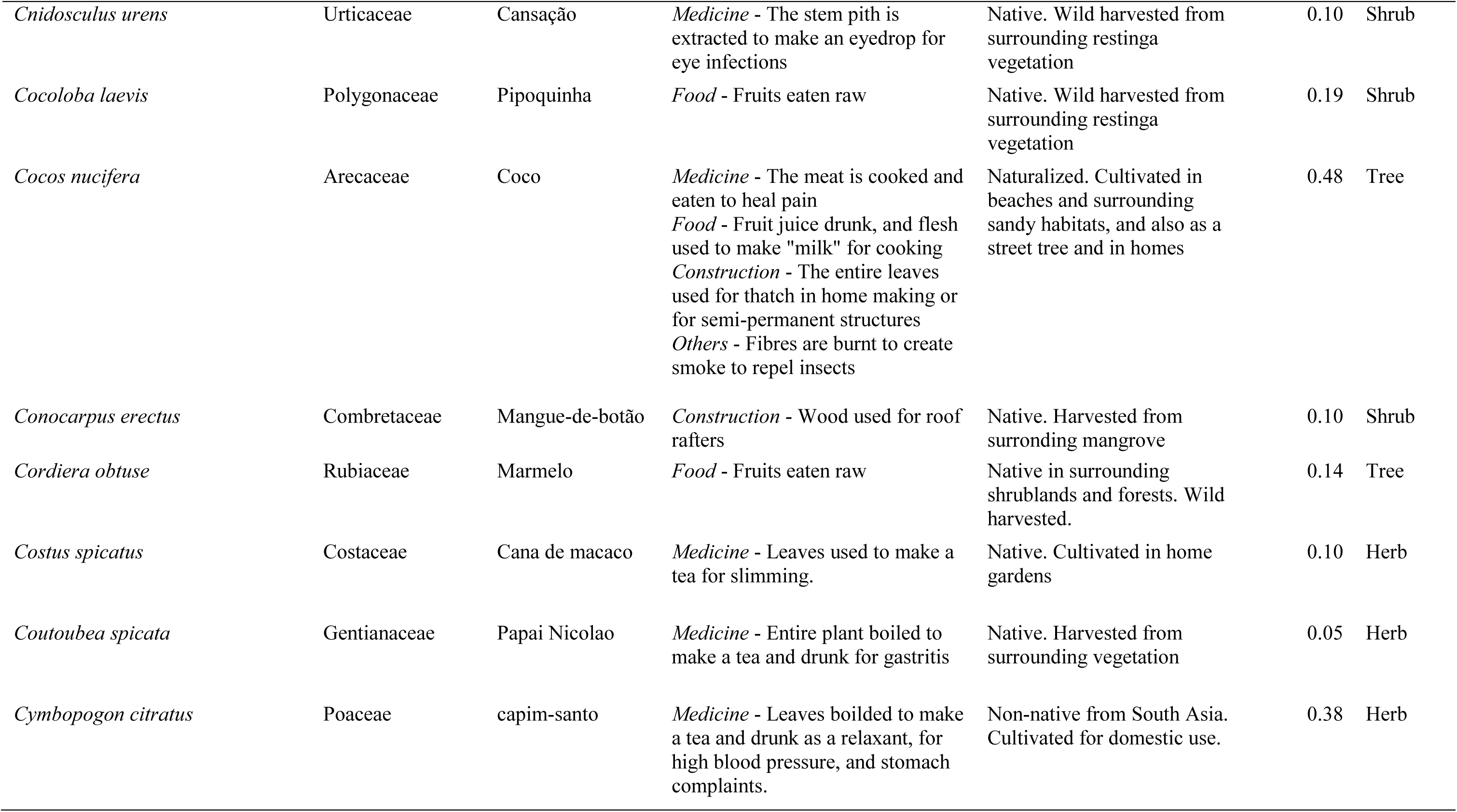
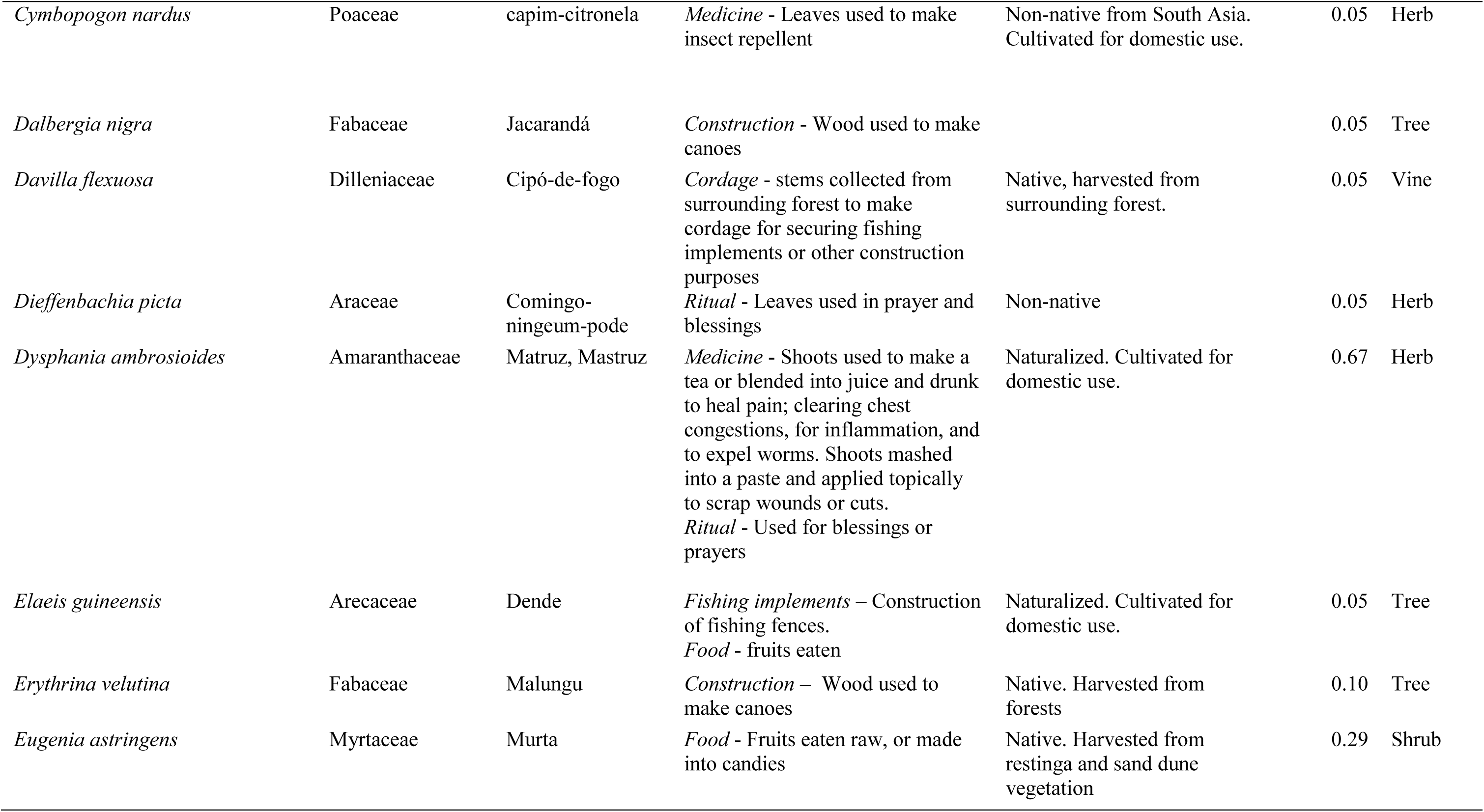
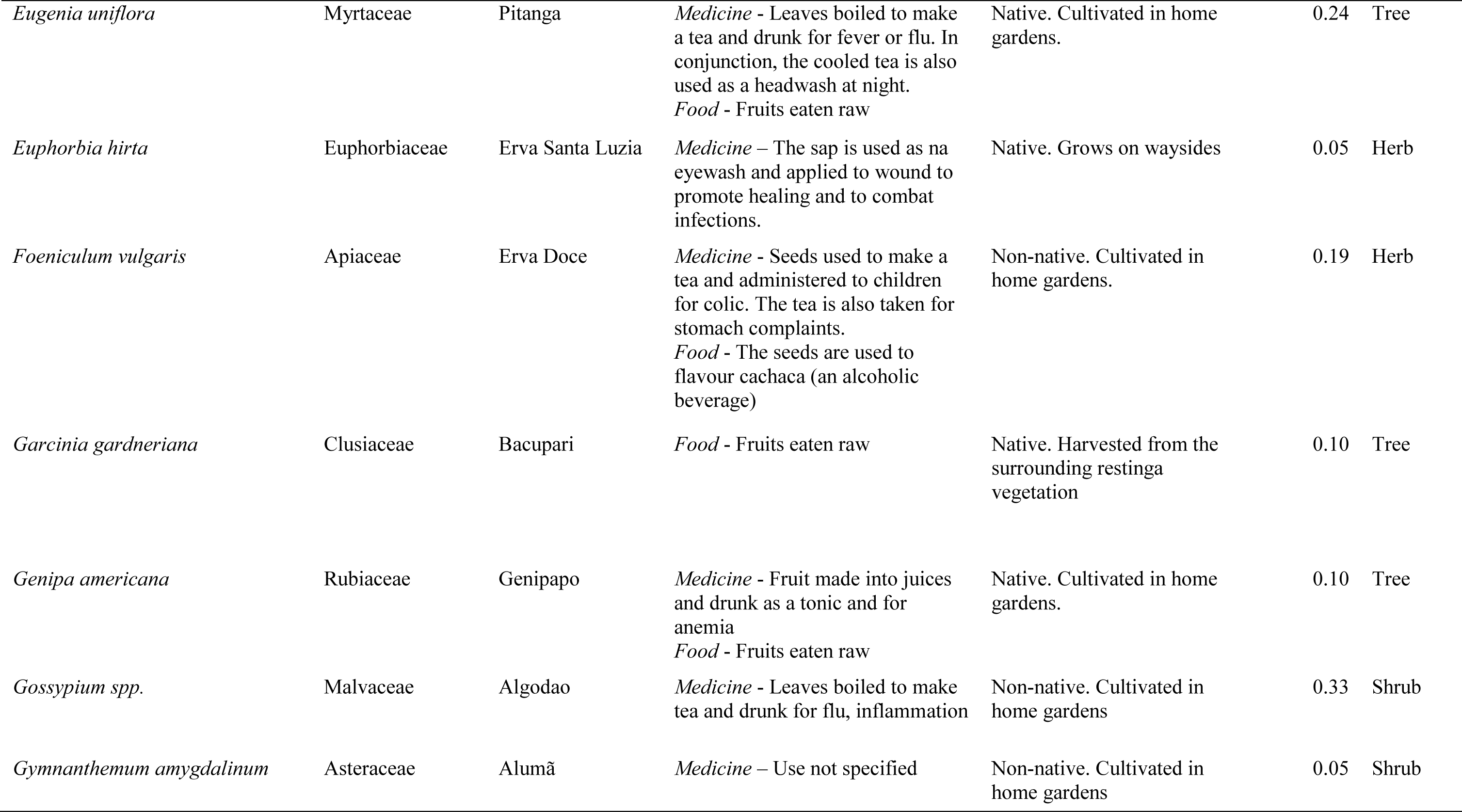
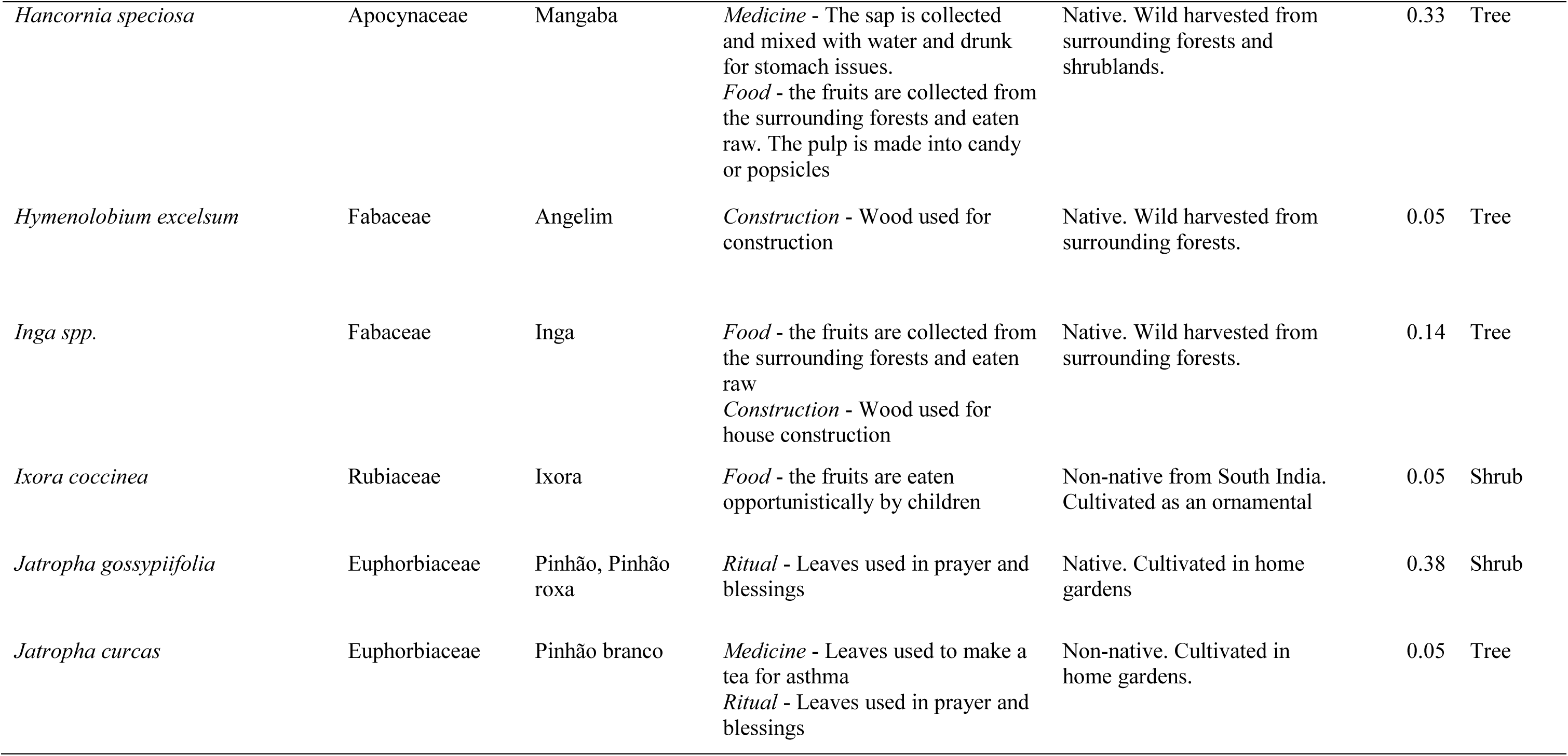
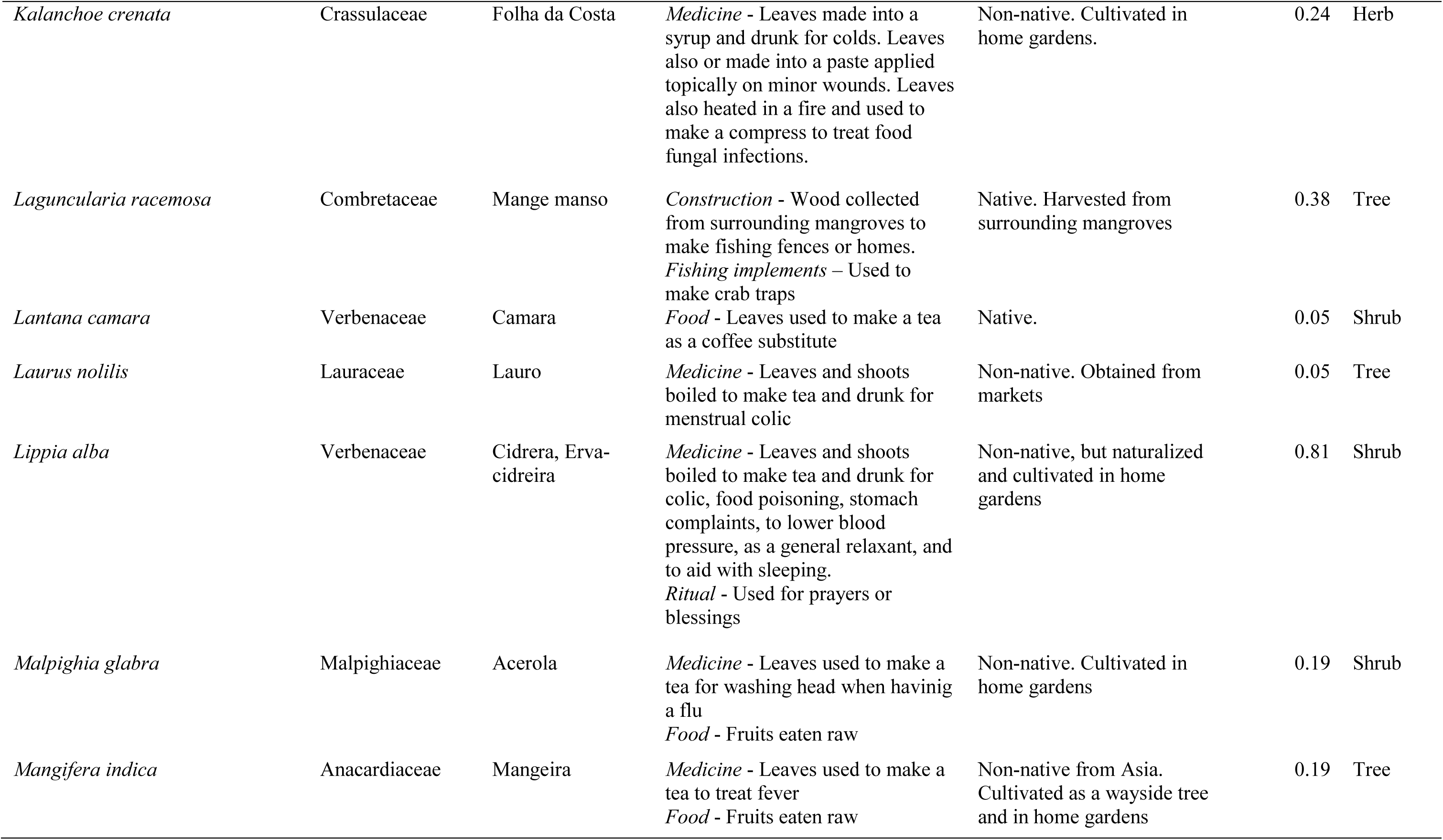
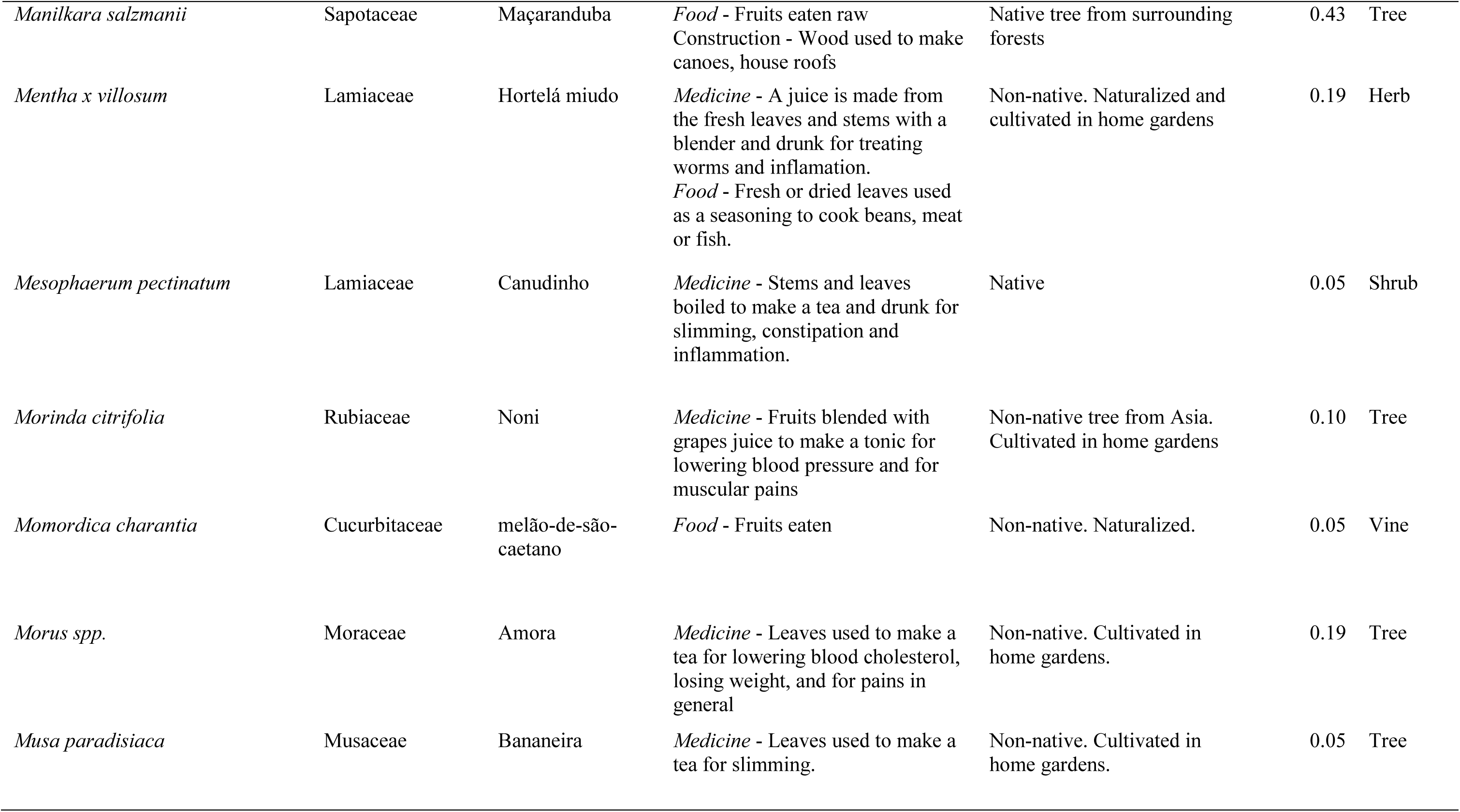
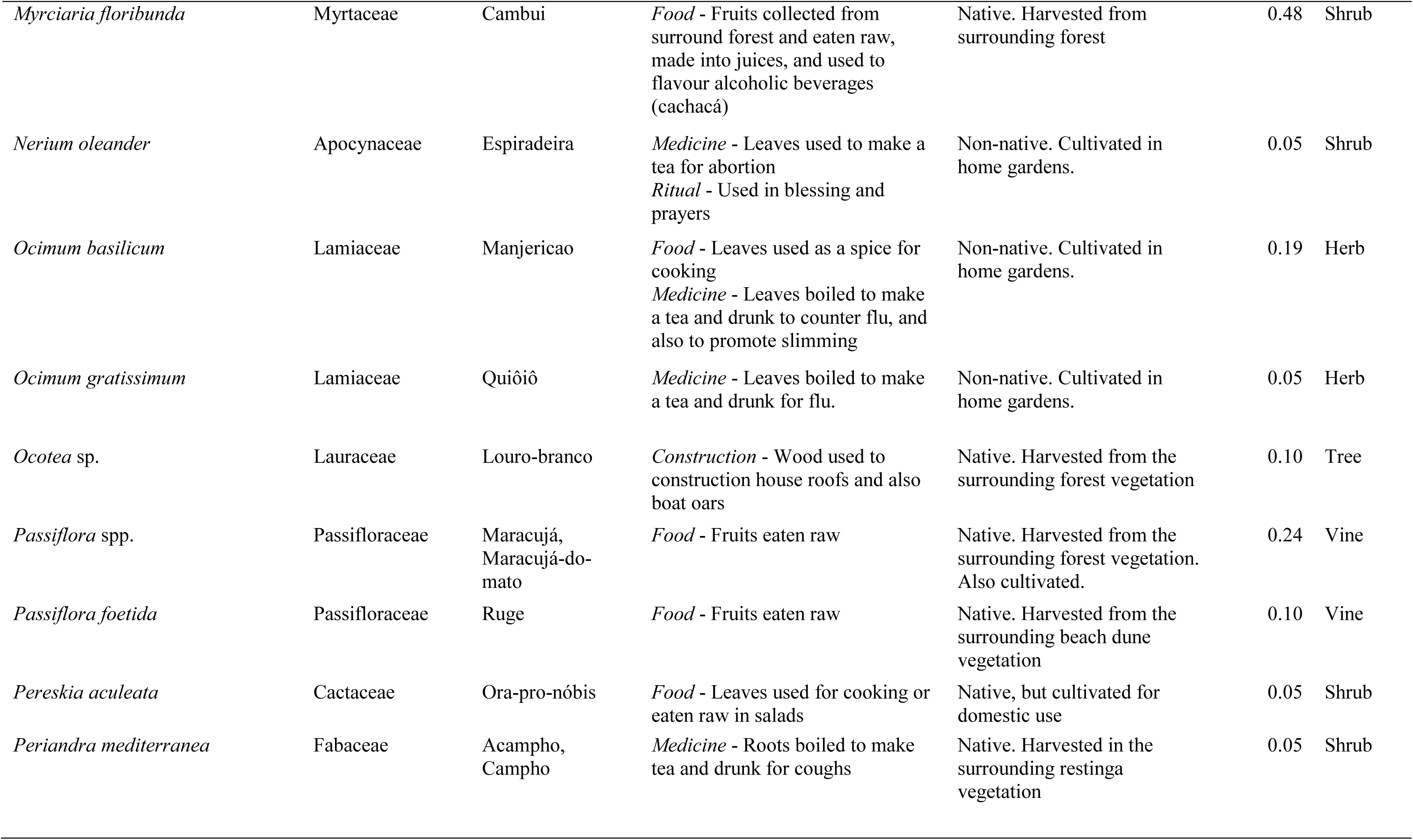
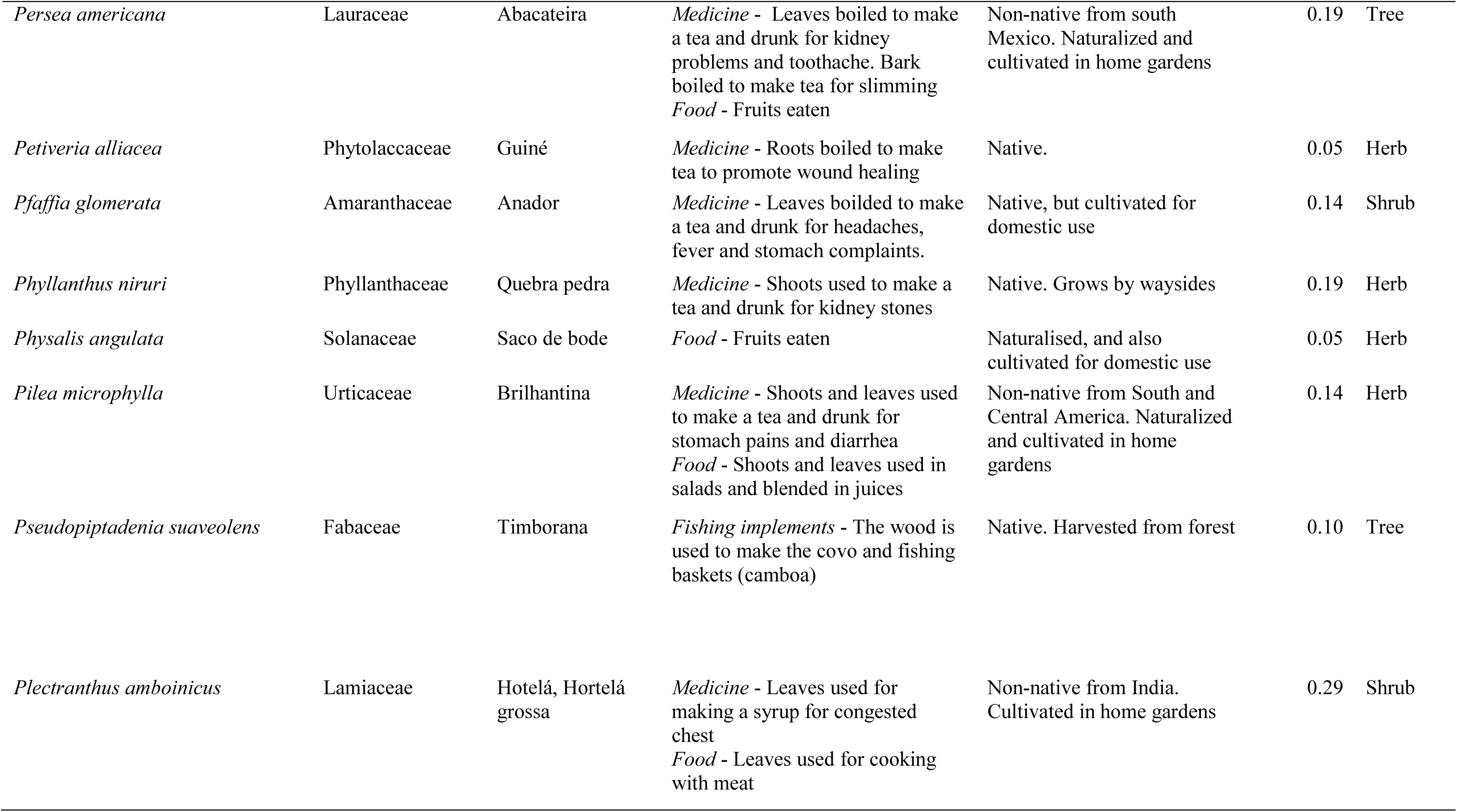
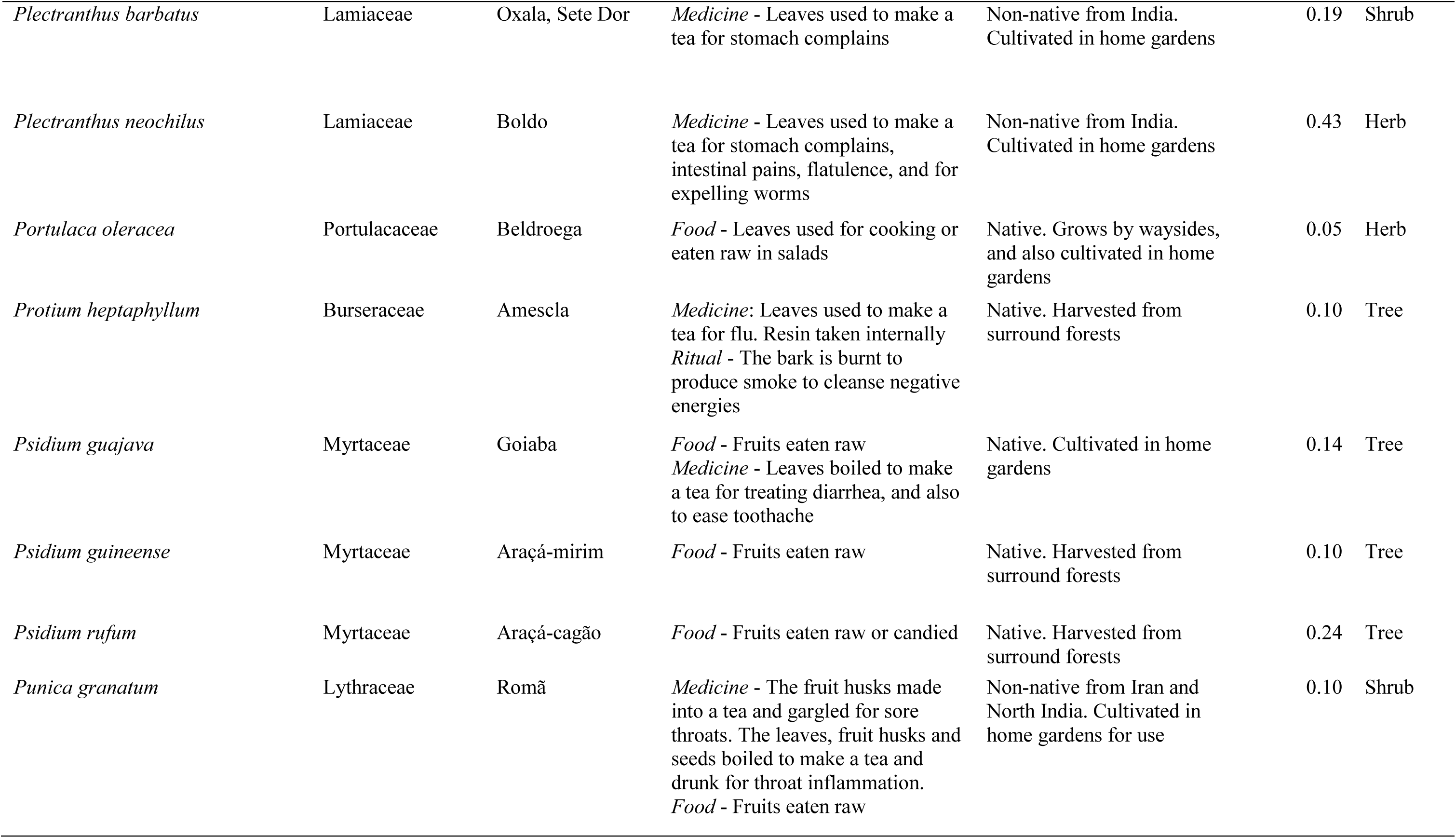
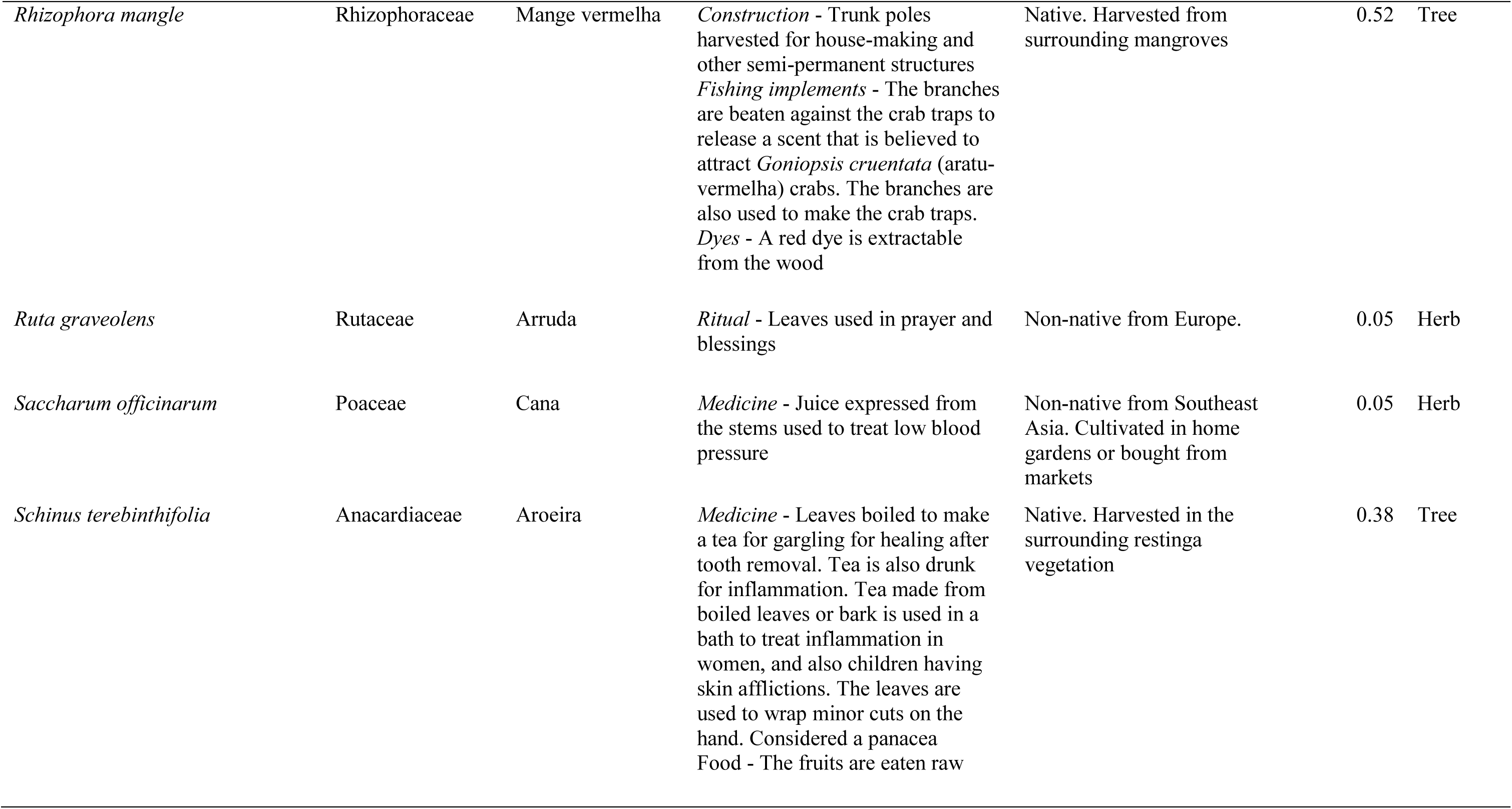
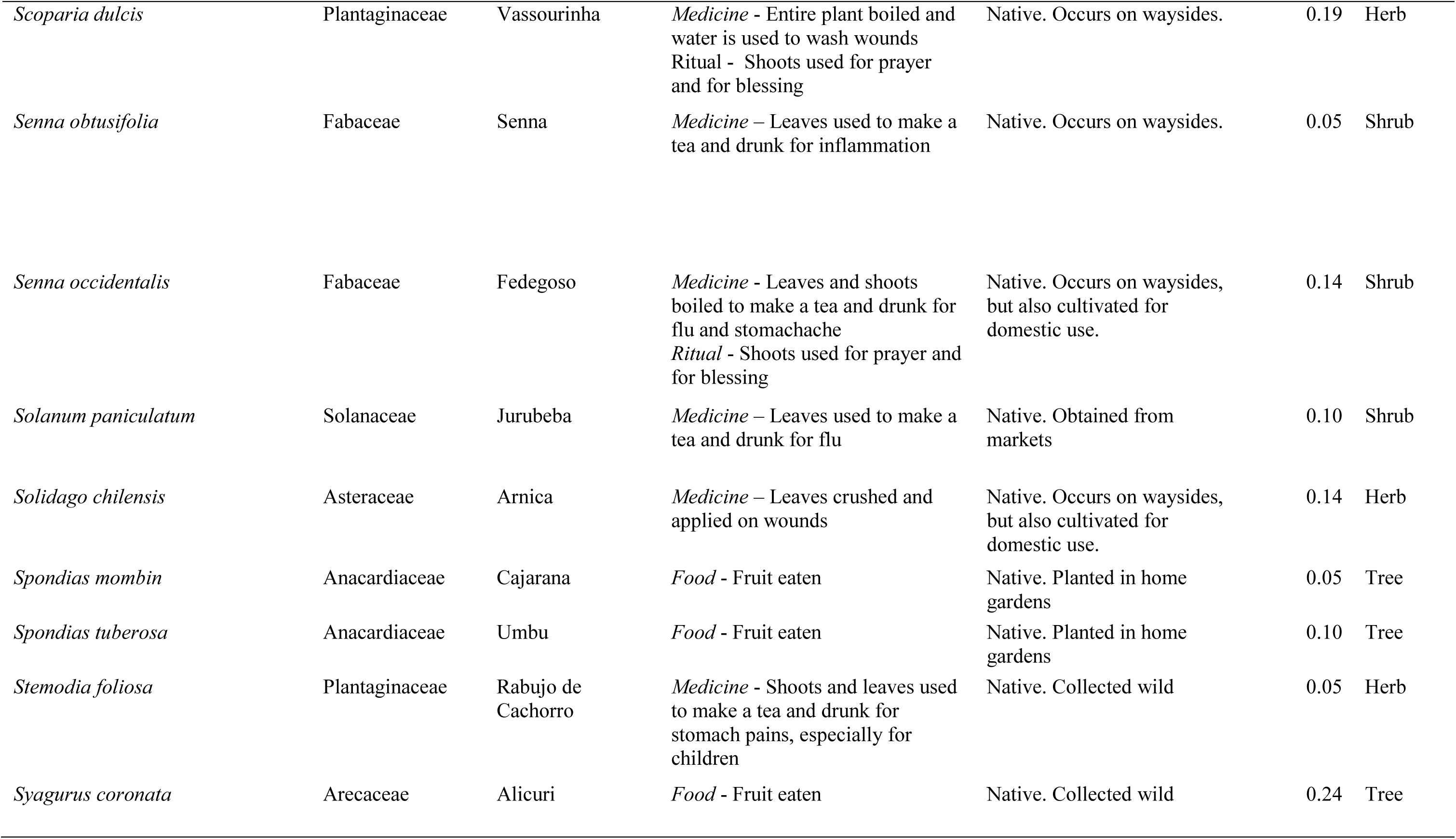
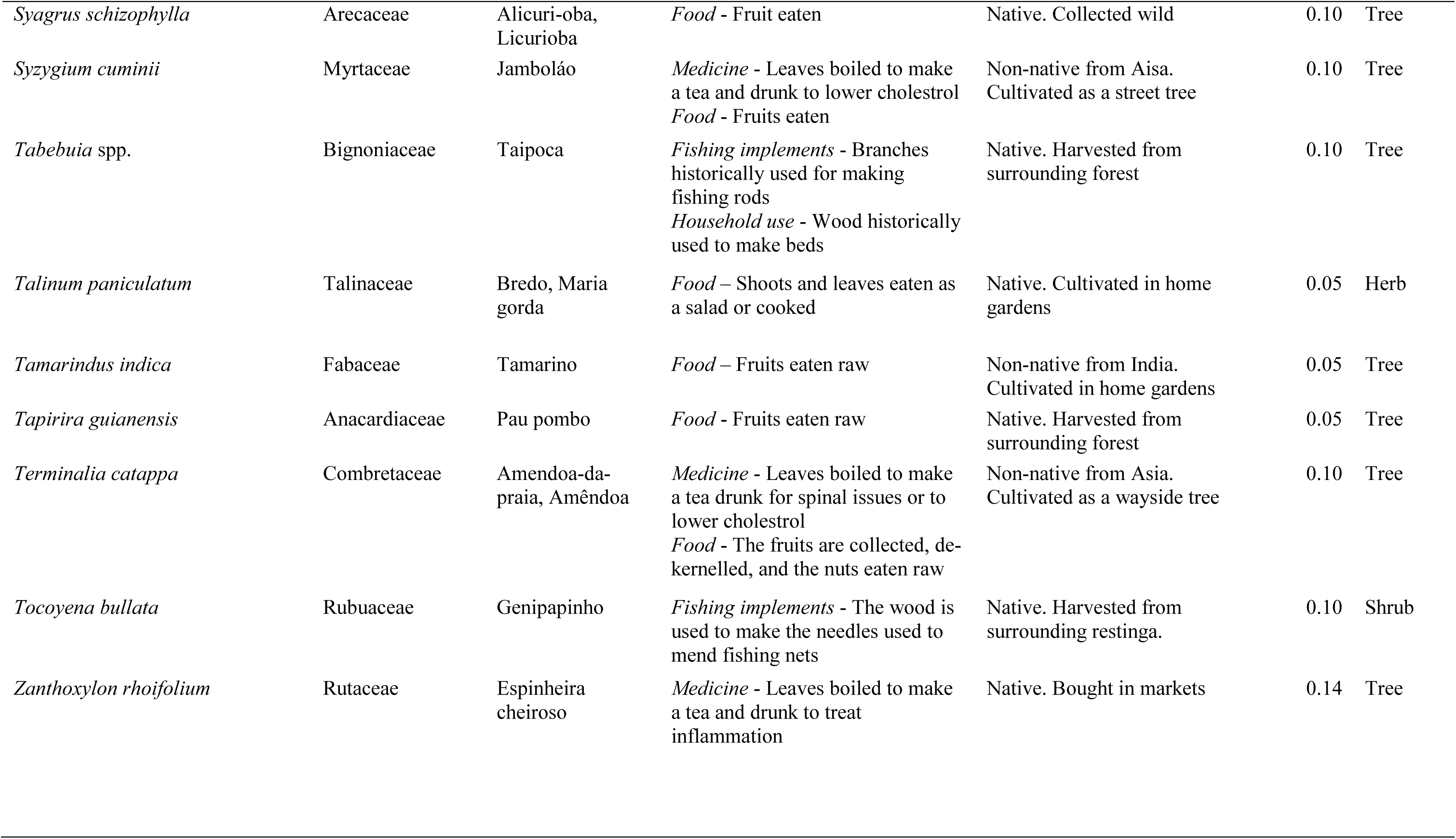
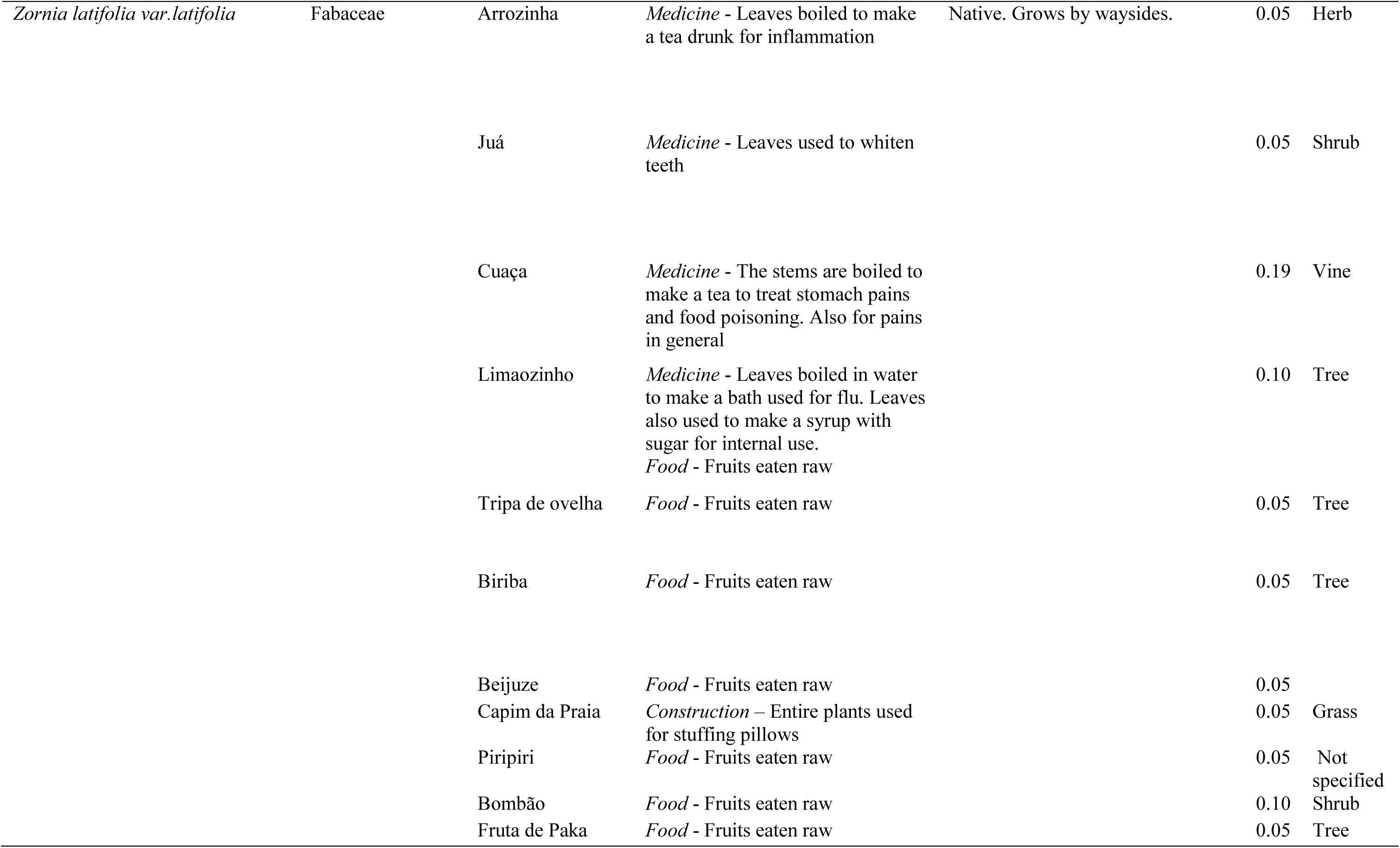
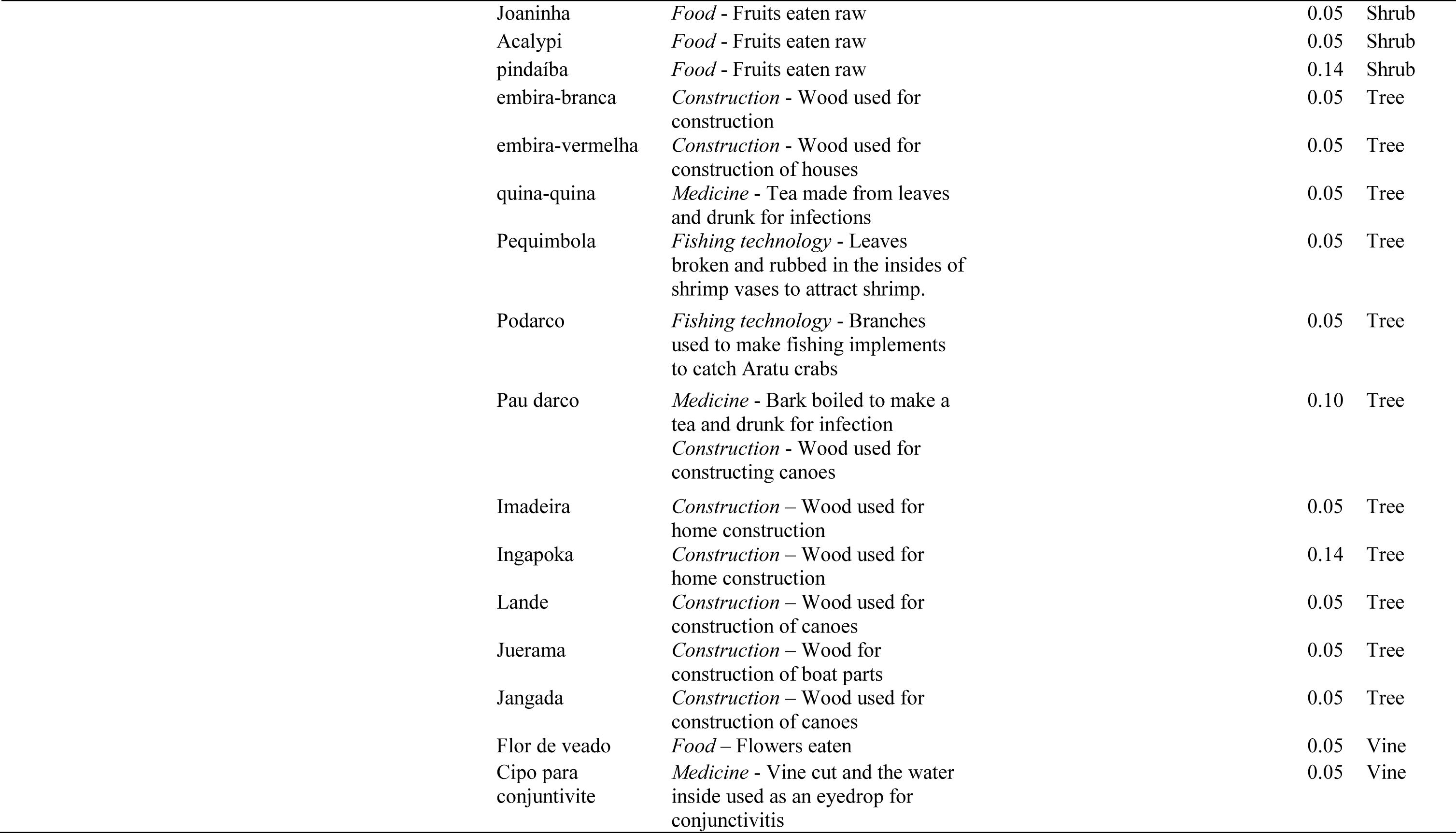
Traditional plants use in artisanal fishing communities in northeast Bahia, Brazil.

Medicine and food plants comprised the highest number of citations (72 and 69 spp. respectively), and among the food category, 48 spp. may be considered non-conventional plant food. For general construction and technology, traditional experts cited 31 spp., among which 25 spp. were cited for fishing technology. Plants used in religious or mystical contexts numbered at 11 spp. A number of species featured in multiple use categories (Fig. 2a), with the highest number of plants shared between the food and medicine categories (21 spp.).

**Fig. 1.**
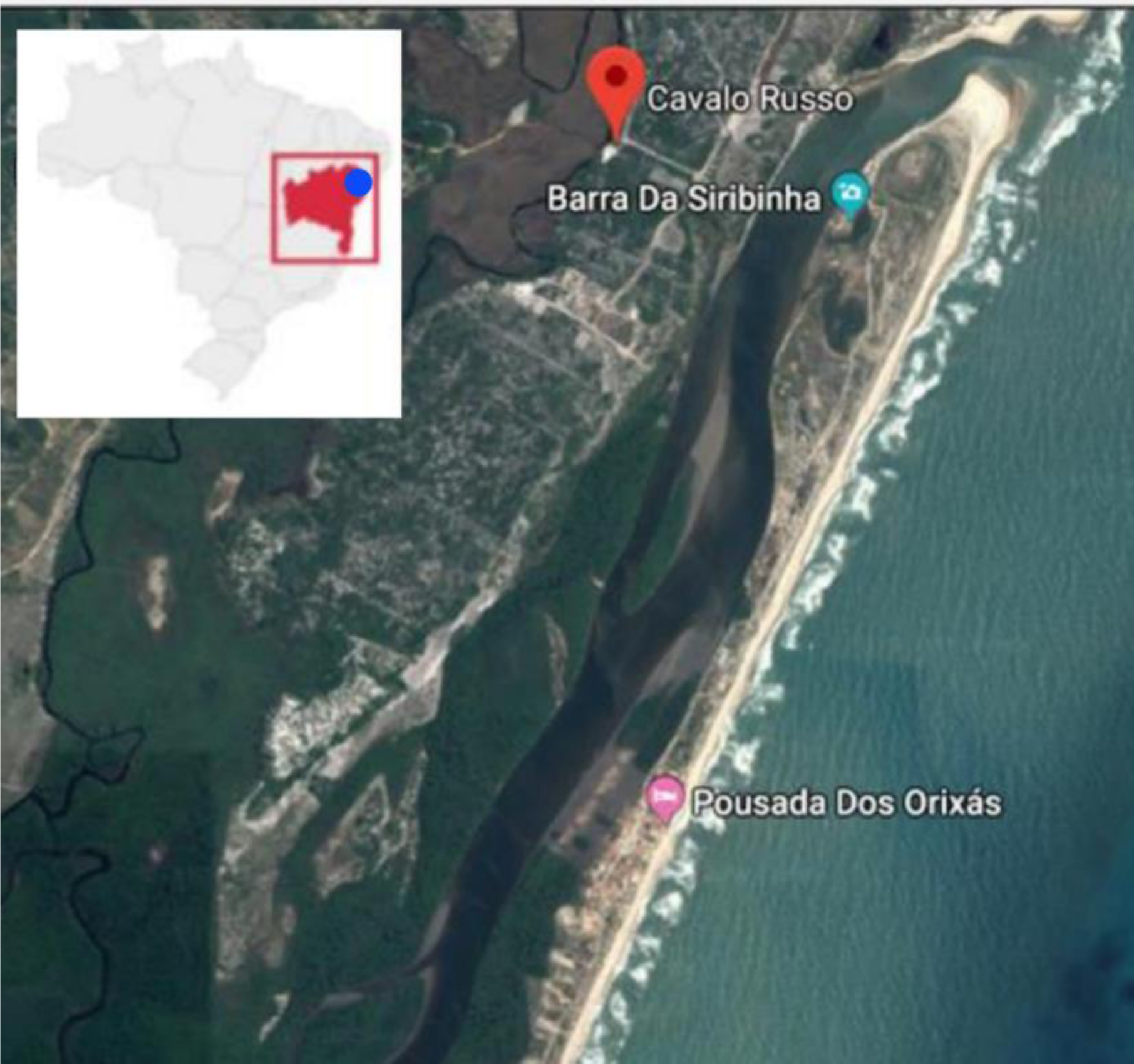
Map showing the locality of the traditional fishermen community at Siribinha, Municipal of Conde, Bahia, Brazil.

**Fig. 2.**
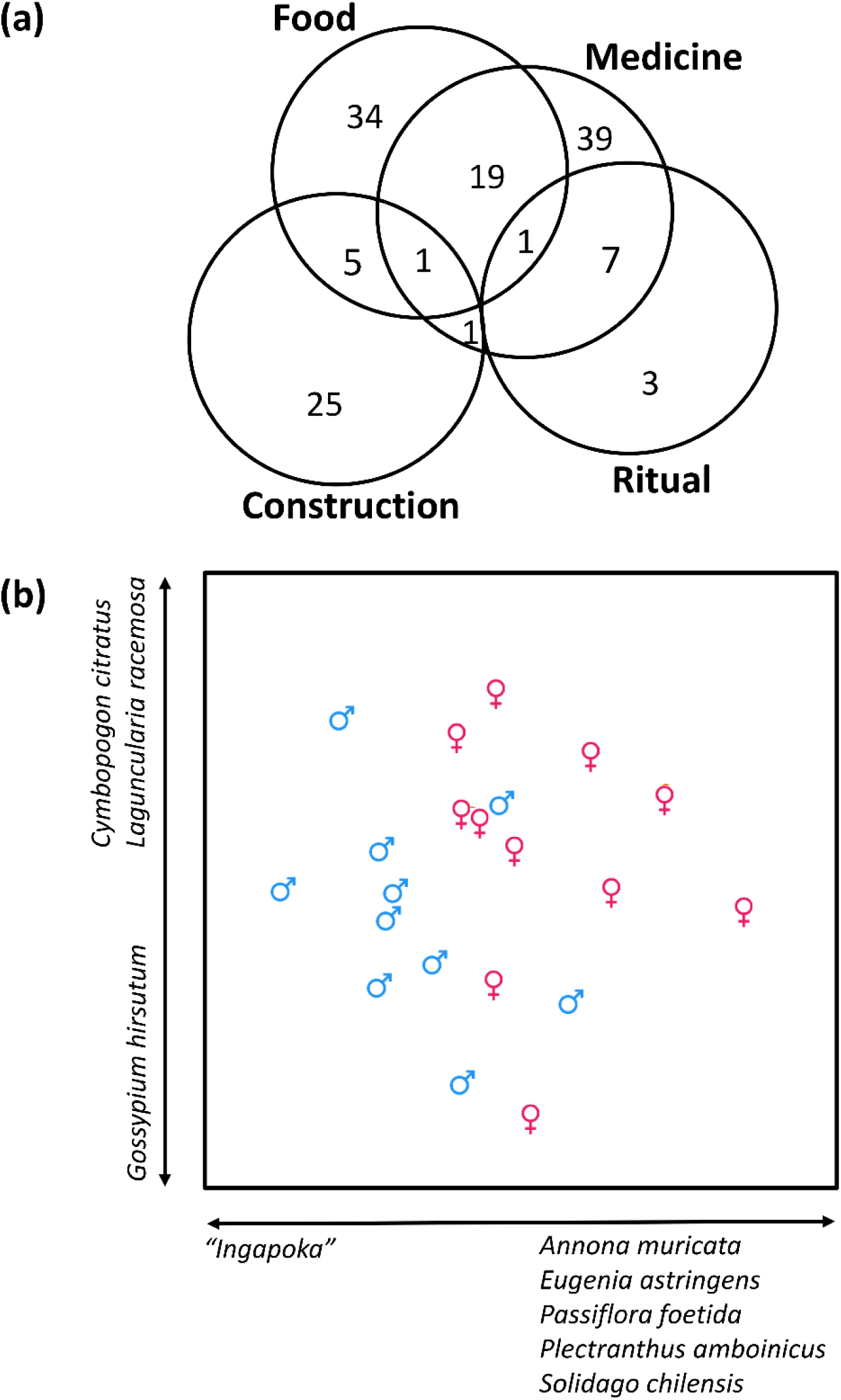
Venn diagram (a) showing the number of plants cited within each plant use category. Note that there were no species which were cited which had both ritual and construction purposes, or both food and ritual purposes. The patterns of plant knowledge of 21 traditional experts (depicted by male and female symbol marker) is visualized in a Non-metric multidimensional scaling ordination, which shows a gender segregation in terms of the plants cited. The plants listed along each axis represent the species which had a Spearman-rank correlation of r > 0.5 or < −0.5 with each ordination axis, signifying their influence in the ordination patterns.

In general, we found high information consensus values (ICF) within all categories of plant use, distribution type and lifeform (all ICF ≥ 0.65), with the highest ICF values documented in plants used for religious or mystical purposes (Table 2). Female traditional experts in general cited on average more plants (24.7 spp.) than male traditional experts (15.9 spp.), although this was not significant. However, female traditional experts cited significantly more medicinal plants, food plants, non-native species and plants from the herb lifeform than males.

**Table 2.** Mean number of citations of plant uses within each category by male and female traditional experts in a traditional fishermen community in northeast Brazil. ^#^Note that numbers for the plant distribution category are calculated based only on the 116 plant species which we could identify at least to genus level. Note also that we did not calculate means and inferential statistics for climbers because of the low number of citations in this category. T-statistics of t-tests and P values comparing the number of citations of plants under each category by females and males and are presented. Significant P values are highlighted. The Informant Consensus Factor (ICF) denotes the degree to which experts within the community may exchange information about plant use, where high values (nearing 1) mean that the plant use information within each respective category is well exchanged among experts, and low values denoting the contrary.

|  | All informants | Females | Males | T (p) | ICF |
| --- | --- | --- | --- | --- | --- |
| All | 20.5 | 24.7 | 15.9 | 2.034 (0.056) | N.A |
| Medicinal | 11.9 | 15.6 | 7.8 | **3.365 (0.003)** | 0.72 |
| Food – All | 10.9 | 13.9 | 7.6 | **2.316 (0.032)** | 0.70 |
| Food – non-conventional<br>plant foods only | 7.4 | 8.8 | 5.8 | 1595 (0.127) | 0.69 |
| Construction &<br>technology in general | 4.6 | 3.9 | 5.3 | 0.885 (0.391) | 0.68 |
| Fishing technology only | 3.3 | 2.7 | 3.9 | 1.086 (0.293) | 0.65 |
| Religious/Mystical | 2.5 | 2.8 | 2.2 | 0.549 (0.590) | 0.81 |
| *<sup>#</sup>Plant distribution or*<br>*lifeform* |  |  |  |  |  |
| Native species | 10.1 | 11.5 | 8.5 | 1.057 (0.304) | 0.69 |
| Non-native species | 7.8 | 10.9 | 4.3 | **4.860 (> 0.001)** | 0.70 |
| Tree | 8.6 | 10 | 7.1 | 1.406 (0.176) | 0.73 |
| Shrub | 5.5 | 6.6 | 4.3 | 1.470 (0.157) | 0.73 |
| Herb | 3.9 | 6.0 | 1.4 | **4.689 (> 0.001)** | 0.65 |
| Vine | N.A | N.A | N.A | N.A | 0.73 |

In corroboration, male and female traditional experts appeared to have different combinations of plant knowledge, with female traditional experts citing a group of five species used variously for food or home grown remedies which were not cited by males, while males cited a species used for construction that were not cited by females (Fig. 2b). Age of the traditional experts did not have a significant bearing on the number of plants each traditional experts cited (Linear regression, R^2^ = 0.02, p = 0.53).

### Medicinal plant use

The most widely reported plants used for medicine were *Cymbopogon citratus, Dysphania ambrosioides, Lippia alba* (Fig. 3a), *Pimpinella anisum*, and *Plectranthus neochilus*. Of the 72 species of plants used for medicinal purposes, 42 species or 58% are commonly cultivated in home gardens for use. However, a number of plants native to the surrounding forest and sand dune vegetation are also collected for medicinal purposes, namely *Aristolochia labiata, Chrysobalanus icaco* (Fig. 3b), *Periandra mediterranea*, and *Schinus terebinthifolius,* and *Stemodia foliosa*.

**Figure 3.**
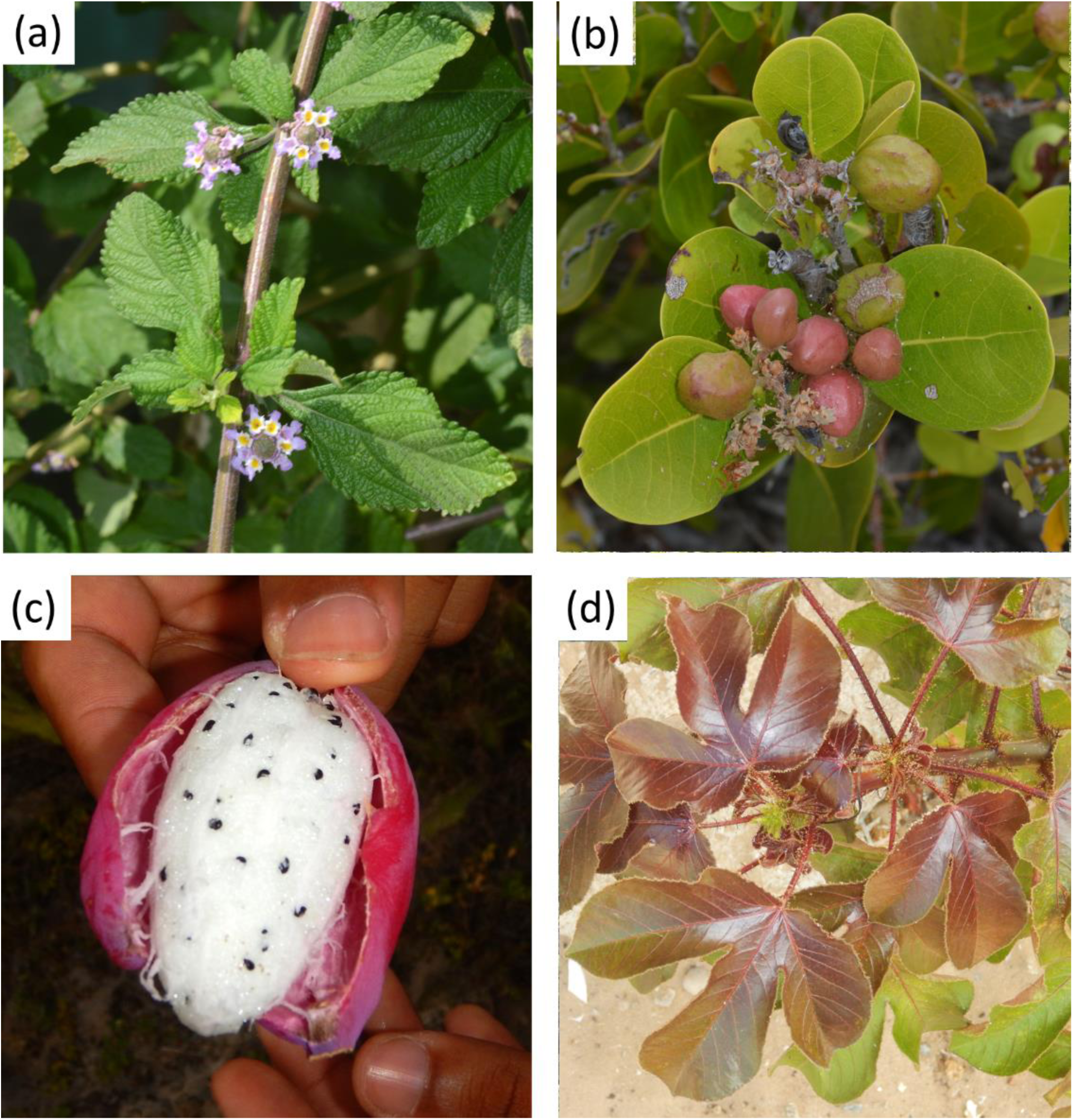
Medicinal plant use in traditional fishermen communities in the littoral north of Bahia, Brazil includes cultivated plants used broadly in the region such as (a) *Lippia alba* (Verbenaceae) and also (b) wild-harvested species such as *Chrysobalanus icaco* (Chrysobalanaceae). Non-conventional plant foods such as (c) *Cereus fernambucensis* (Cactaceae) are also utilized by the community. The leaves of (d) *Jatropha gossypiifolia* (Euphorbiaceae) are used in blessing rituals.

The most common method of medicinal plant use within the community is by the preparation of teas from leaves, stems or roots (Table 1). A number of plants are also made into juices or into syrups in conjunction with other ingredients. An interesting mode of using a medicinal plant, *Hancornia speciosa*, involved stirring the raw leaf sap into water and drinking the concoction. Some plants are also processed into a paste and applied topically to wounds (e.g. *Dysphania ambrosioides*), or shoots boiled in water and used for bathing (e.g. *Anacardium occidentale* and *Schinus terebinthifolius*) or for washing wounds.

### Uses of non-conventional edible plants and other non-conventional plant foods

The utilization of seasonal bush fruits from the surrounding forests and *restinga* shrublands continues to be widely practiced, and traditional experts report the harvesting the fruits and nuts of the native palms (primarily *Allagoptera brevicalyx*. *Bactris* spp., and *Syagrus* spp.), cashew nut tree (*Anarcardium occidentale*) and also the fruits of various members of the Annonaceae, Cactaceae (Fig. 3c), Clusiaceae, Myrtaceae, Polygnonceae, and Sapotaceae (Table 1).

However certain specific uses of some species are no longer practiced, such as the extraction of milk from the inner flesh of *Syagrus* spp. and *Allagoptera brevicalyx* fruits.

### Plants used for general construction, fishing technology and daily-use implements

The wood of *Rhizophora mangle* was the primary species used for house making in the past (Table 1), although most homes in the community are now made of concrete. Other species of trees were also used in construction or to fortify the structure of houses (e.g. *Laguncularia racemosa, Hymenolobium excelsum*, etc.) (Table 1).

A number tree species was cited as being used for fishing technology. Fishing canoes were made using wood of tree species such as *Artocarpus heterophyllus, Dalbergia nigra, Erythrina velutina*, etc., although *A. heterophyllus* is held in high esteem for this purpose. The leaf stalks of palm species *Attalea funifera, Cocos nucifera*, and *Elaeis guineensis* were also sought-after for making hatch for houses, and also boat shelters (Fig. 4a). Additionally, *Attalea funifera*, and *Elaeis guineensis* are used to make artisanal fishing fences (*camboas*) and crab traps (*covos*) (Figure 4b). These leaf stalks were processed into strips, shaped and bound by cordage obtained from plants such as *Davilla flexuosa*. Another notable example of fishing technology includes the use of *L. racemosa* branches beaten against the crab traps, which is believed to exude a scent attract crabs.

**Figure 4.**
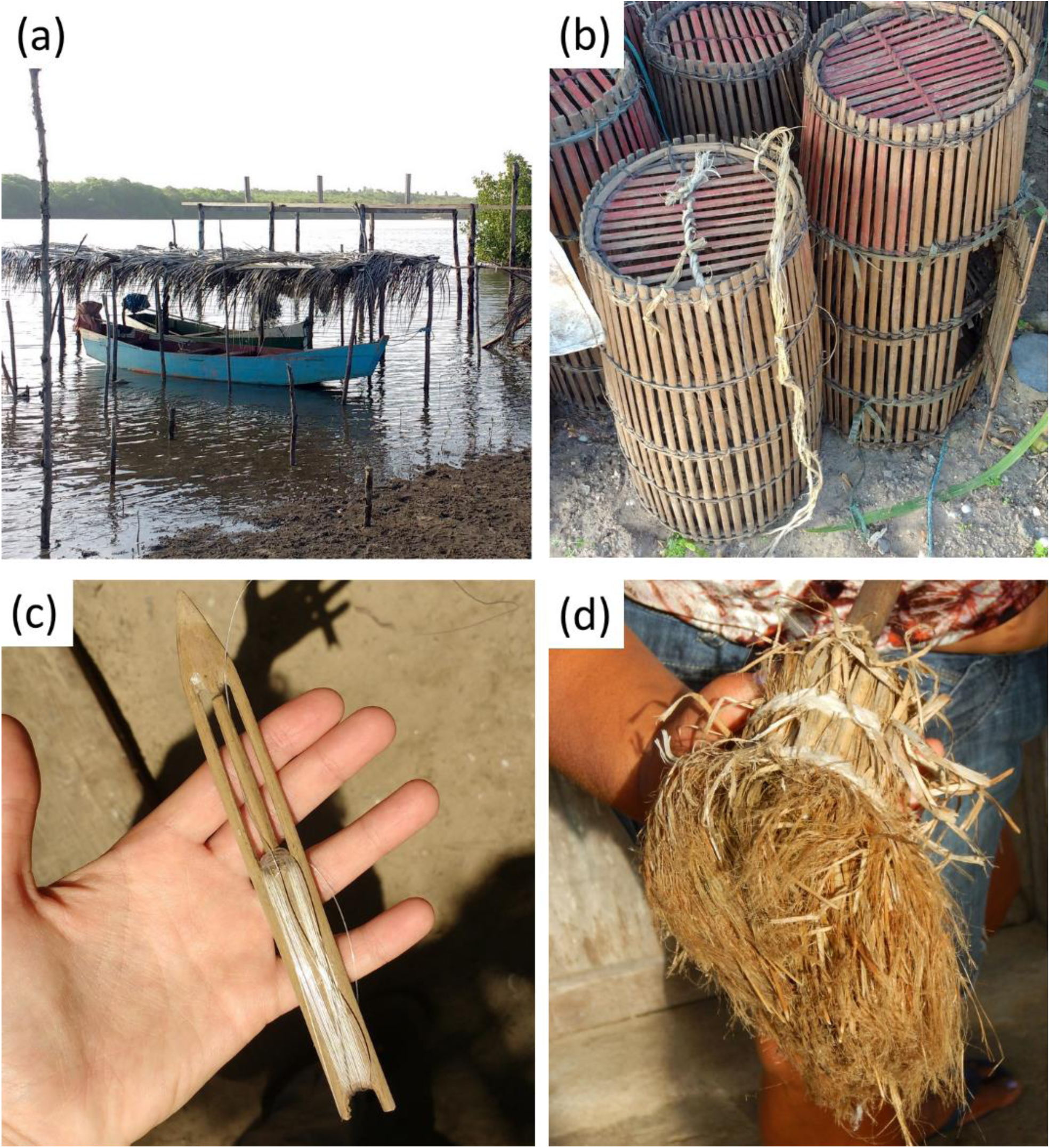
Wood resources and fibres used in construction, fishing practices and daily-use implements in a artisanal fishing community in the littoral north of Bahia, Brazil. The wood of mangrove tree, *Rhizophora mangle* (Rhizophoraceae) and palm leaf thatch used to construct (a) boat shelters; (b) slivers of the rachis of the palm *Attalaea funifera* (Arecaceae) bound together by cordage of *Davilla flexuosa* (Dilleniaceae) to make crab traps; (c) a net-mending needle carved from *Tocoyena bullata* (Rubiaceae); broom heads made from the leaf fibres of *A. funifera*.

Before nylon nets were widely available, the community relied on plant fibres to weave nets. Of the respondents interviewed, many mentioned the use of the leaf stalks of the native palm *Bactris setosa* (Arecaceae). The women collected the leaves and did the initial processing (beating the leaf stalks to obtain the fibres), while the men hand-wove nets using a spool. The process of net-making was labour intensive, and is a process that has been discontinued in favor of widely available nylon nets. Additional fishing implements constructed from locally-sourced plant material included needles for net mending (Figure 4c), for which the wood of *Tocoyena bullata* was used.

Palm leaves were also historically used to transport of fish out of the community to regional markets, whereby fishermen wove the leaflets of the *Syagurus* spp. or *Cocos nucifera* palm fronds through the flesh of the fish to secure them onto carrying poles. Other uses of palm fibre included the use of *Attalea funifera* leaf stalk fibres to make broom heads (Fig. 4d).

### Plants used in spiritual and religious purposes and the role of faith

The catholic and evangelic denomination of Christianity are the main religions practiced within the community, and the community role of a “blesser” or *rezadeiro* is designated to a person who blesses or prays for members of the community upon request. *Rezeiros* used to play a prominent role in the community as the person consulted for blessings and faith healing, but this practice is now restricted to the older generation. A number of plants were listed as being used specifically in praying or bestowing blessings, namely *Dieffenbachia* spp., *Jatropha gossypiifolia* (Fig. 3d), *Scoparia dulcis*, and *Senna occidentalis*. However, according to a *rezadeira* (female *rezadeiro*) we interviewed, the words used are more important than the plants, and she says that any plant will suffice.

Importance is also attributed to the number of branches used in blessing. For bestowing blessings or prayers, the *rezadeira* we interviewed collects three branches and she performs the prayer six times over the recipient, after which she recites another prayer to discard the branches. In such prayers, the negative energies that cause sickness in a person are believed to be transferred to the leaves or branches that are discarded.

Some community elders emphasized also the importance of faith in healing, and attributes huge importance to the recipients of healing having faith in the healing. However, one elderly male traditional expert elaborated on certain ritual practices in his use of medicinal plants whereby branches picked for making tea are collected in odd numbers, principally three or five branches, but never six.

## Discussion

Globally, cultural knowledge of rural communities is under threat from urbanization, and it is therefore critical to register and understanding patterns of this cultural knowledge for future generations. Studying artisanal fishing communities in the littoral north region of Bahia, northeast Brazil, we found a diverse cultural knowledge of plant use for medicinal, construction, technological and ritual purposes. It is clear also that the women of the community in general harbor more plant knowledge than the males.

### The broader context of medicinal plant use

The medicinal plants used by the communities fit within belong a broader regional milieu of medicinal plant knowledge (di Stasi et al. 2002; Lorenzi & Matos 2002), and the domestic cultivation of the most commonly cited species *Anarcardium occidentale, Cymbopogon citratus, Dysphania ambrosioides*, and *Lippia alba* (Fig. 3a) have variously been reported for other indigenous Indian communities in the region (Alberquerque et al. 2009; Borges & Bautista 2010; Cunha et al. 2012) and afro-brazilian (Quilombola) communities (Gomes and Bandeira 2012; Mota and Dias 2016; Lisboa et al. 2017) and rural communities (Almeida et al. 2014; Bandeira et al. 2015). Many of the species cited are also available commercially in regional markets within the state (de Araújo et al. 2018; pers obs). Likewise, the use of various species of *Ocimum* as reported in this study has also been previously documented in the region (de Holanda and de Albuquerque 1998)

However, some members of the communities are also stewards of knowledge regarding medicinal plants in the natural vegetation in the area. The use of *Anacardium occidentale* and *Schinus terebinthifolia* for skin problems and female genitourinary disorders are in line with recent reports for other riverine fishermen communities in the region (Paiva et al. 2017). Other notable examples are the wild harvested *Chrysobalanus icaco* (Fig. 3b) and *Periandra mediterranea*.

The common names applied to medicinal plants deserve special mention. In one particular instant, we came across a common name, *acamfo*, applied to *Periandra mediterranea*, for which could not find references to in the literature. Previously, other workers had reported the use of this plant under the names *acançu* (Almeida et al. 2010), *arcancuz* (da Silva et al. 2012), *alcacuz* and *alcancuz* (Gomes et al. 2012) from various other localities inland in the state. This site specific differences in names may reflect phonetic changes in a local name being applied to a single plant, as people from communities move from place to place. Sematic differences in common names can sometimes also reflect taxonomic differences, such as in the case of the genus *Passiflora* (Neto 2008). On the other hand, some local names refer to different species depending on region. In our study region, *carqueja* is applied to *Borreria verticillata* (Rubiaceae), but south in the state of Minas Gerais, *carqueja* refers to species of *Baccharis* (Asteraceae). In both regions, the respective carqueja species are used for treating stomach problems (Lorenzi and Matos 2008), reflecting a convergence in use. The local specialists likely acquired a broad knowledge of medicinal plants by way of experimentation and trading information with relatives or people from the surrounding communities. This appears to be supported by the high informant consensus factor that we obtained for the medicinal plant category. Yet it is also of note that many plants listed by community specialists are herbaceous or naturalized plants, a pattern that has been noted by Voeks (1996).

### Food plants

As with medicinal plant use, the use of plants for consumption as food fit into the two categories of cultivated for domestic use and wild harvested (Fig. 3c), and some of the species overlap with plants used for medicine. The knowledge on the use of wild plants appear to be passed on to the younger generation, and children from the community continue to harvest seasonal fruits for consumption (Fig 3d; pers. ob.), and this is important avenue of follow up research. Various of the plants used have also been reported for other communities in the Bahia state (Rodrigues and Guedes 2006; Agra et al. 2008; Neto et al. 2014; do Nascimento et al. 2015; Mota and Dias 2016).

### Plants for construction and technological purposes

The plants used for construction and technology have been documented by various workers to be used in the region. For instance, leaves and fibres of palms such as *Attalaea funifera, Bactris* spp. and *Syagrus* spp. feature prominently in traditional use in the region (Lorenzi et al. 2000), and the use of mangrove wood to construct houses was a matter of convenience for communities living near mangrove environments (Bezerra 2008). However, in the present, wood material for construction of houses and particularly boats are largely sourced from eucalypt plantations, and cordage have largely been replaced by commercial nylon.

Canoes were previously constructed using a single trunk, but later wood planks were used. Wood used for the construction of canoes presents a divergence from previous literature (Andrade et al. 2016). Among the species cited in Andrade et al. (2016) the current study only had *Artocarpus heterophyllus* in common.

It is interesting to note that in other ethnobotanical studies on other littoral communities, the use of species for constructing fishing technology was not mentioned (Lopes et al. 2013).

### Religious and mystical use of plants

We describe for the first time specific details and attitudes relating to religious and ritualistic practices involving the use of plants in blessing and prayers. Notably, most of the plants listed under this purpose are naturalized or cultivated, with the exception of *Protium heptaphyllum* and *Schinus terebinthifolius*, and the species cited accord with those reported by other authors (Varella 1973; de Oliveira et al. 2009). Although members of the communities are predominantly Christian, the influences of the syncretic African Candomblé and Umbanda religions is conspicuous, particularly in the use of *Jatropha gossypiifolia* (Varella 1973; Pires et al. 2009).

### The maintenance or erosion of traditional knowledge

Although the majority of plant medicinal uses is well documented, there was a consensus among the local specialists that this knowledge is undergoing a progressive dilution with the proceeding generations. Nevertheless, we still know little about the degree of knowledge transfer across age groups within the community, as we only interviewed traditional experts aged 43 and above.

The paradigm of loss of cultural knowledge or that succeeding generations have less knowledge due to globalization has been challenged on at least one count by Vandebroek and Balick (2012), who studied people who use plants to self-medicate in the Dominican Republic. Indeed, informal interactions with children in the community gave us the impression that the younger generation do have some traditional knowledge about food and medicinal plant uses, and more formal interviews with people of all age groups in the community would be a worth follow up study.

More critically, the landscape change in the region has meant that certain forests where medicinal plants used to be collected have now been felled. As has been reported from other communities in the state of Bahia, the process of modernization, particularly increasing access to formal education and availability of modern pharmaceuticals, may lead to traditional domains of medical knowledge being increasingly perceived as an irrelevant province of past generations (Voeks & Leony 2004). Indeed, environmental degradation and large changes in modern social and economic systems are well known to be drivers leading to a reduction in traditional medicinal plant use (Anyinam 1995; Srithi et al. 2009). While we do not quantitative figures, it surfaced during interviews with some of the traditional experts that certain medicinal plants used to be collected from some forests that are no longer extant.

The substitution of artificial material used in fishing technology is probably one of the more prominent observations made during the study. As far as we could ascertain from informal queries and observations, there are no longer any nets made of plant material due to the availability of more durable nylon thread, and the knowledge of how to process the plant fibres to make such nets are only known by the elderly generation. To preserve this aspect of artisanal knowledge, videos are being recorded of how fishermen used to make nets, and also museum artifacts are being gathered.

## Conclusion

The registration of a rich tradition of plant use by a fishing community is surprising, and underpins an urgent need to conserve traditional knowledge in the region. Educational programs for elementary school students involving the dissemination of this body of knowledge traditional plant use knowledge are in the works, with hopes that the next generation will become custodians of their cultural knowledge. Further studies are needed to document plant use of other traditional communities within the river system.

## Acknowledgements

We thank Domingos Cardoso and Maria Lenise Silva Guedes for help with identifying some specimens. We thank also the member of the communities who participated in interviews. The study was supported by a INCT grant awarded to Charbel El-Hani.

